# Gene-environment correlation: The role of family environment in academic development

**DOI:** 10.1101/2024.01.05.574339

**Authors:** Quan Zhou, Agnieszka Gidziela, Andrea G. Allegrini, Rosa Cheesman, Jasmin Wertz, Jessye Maxwell, Robert Plomin, Kaili Rimfeld, Margherita Malanchini

## Abstract

Academic achievement is partly heritable and highly polygenic. However, genetic effects on academic achievement are not independent of environmental processes. We investigated whether aspects of the family environment mediated genetic effects on academic achievement across development. Our sample included 5,151 children who participated in the Twins Early Development Study, as well as their parents and teachers. Data on academic achievement and family environments (parenting, home environments, and geocoded indices of neighbourhood characteristics) were available at ages 7, 9, 12 and 16. We computed educational attainment polygenic scores (PGS), and further separated genetic effects into cognitive and noncognitive PGS. Three core findings emerged. First, aspects of the family environment, but not the wider neighbourhood context, consistently mediated the PGS effects on achievement across development –accounting for up to 34.3% of the total effect. Family characteristics mattered beyond socio-economic status. Second, family environments were more robustly linked to noncognitive PGS effects on academic achievement than cognitive PGS effects. Third, when we investigated whether environmental mediation effects could also be observed when considering differences between siblings, adjusting for family fixed effects, we found that environmental mediation was nearly exclusively observed between families. This is consistent with the proposition that family environmental contexts contribute to academic development via passive gene-environment correlation processes, or genetic nurture. Our results show how parents tend to shape environments that foster their children’s academic development partly based on their own genetic disposition, particularly towards noncognitive skills, rather than responding to each child’s genetic disposition.

## Introduction

Academic achievement during childhood and adolescence is associated with a host of positive life outcomes (1), from better physical health and psychological wellbeing (2,3) to higher earnings (4–6). Students differ widely in academic achievement within neighbourhoods, schools, and even classrooms (7–9). These observed differences are partly due to genetic factors. A meta-analysis of twin studies, which estimate genetic and environmental effects on traits by comparing the observed similarity between pairs of identical and fraternal twins, found that genetic differences accounted for 40% of individual differences in educational outcomes (10), a statistic known as heritability.

Evidence for the substantial contribution of genetic variation to differences between students in academic achievement has also emerged from studies that have applied DNA-based methods. Using genome-wide complex trait analysis (GCTA) (11) and linkage disequilibrium score regression (LDSC) (12), these studies found heritability estimates of ∼30% (13–15). A polygenic score (PGS) constructed aggregating findings from large genome-wide association studies (GWAS) of educational attainment (i.e., years of schooling) (16,17) has also been found to explain up to 15% of the variation in academic achievement at the end of compulsory education (10,16, 17).

However, these genetic effects on academic achievement are not independent of environmental processes (20). Children evoke and select environmental experiences partly based on their genetically influenced psychosocial characteristics, two processes that have been labelled evocative and active gene-environment correlation (21). These transactions accumulate over development, particularly as children gain more autonomy to select their own experiences (22,23). Children are also likely to experience environments that correlate with their genetic dispositions simply by growing up with their biological relatives, a phenomenon referred to as passive gene-environment correlation (21).

Parents are likely to shape children’s rearing environments partly in line with their own genetic dispositions (21,24,25). One study found that mothers’ genetic propensity towards education, indexed by their educational attainment PGS, was associated with children’s attainment after accounting for children’s genetic propensity (26). This genetic nurture (27,28) path from mothers’ genetics to children’s attainment was mediated by early family characteristics, specifically by cognitively stimulating parenting (26).

Building on this initial evidence, the present study aims to systematically investigate how family environments mediate PGS effects on academic achievement over compulsory education, from age 7 to 16. We include multiple aspects of the proximal family environment (e.g., socioeconomic status, parenting and chaos at home), which have all been linked to individual differences in education (29–31). In addition, we consider more distal aspects of the family environment by examining the role of neighbourhood contexts, such as neighbourhood health ratings, occupancy ratings and levels of pollution. Children growing up in disadvantaged neighbourhoods have been consistently found to exhibit worse physical and mental health and have poorer educational and economic outcomes if compared to children from more affluent neighbourhoods (32,33). Previous research also found that neighbourhood ecological risk (including deprivation, dilapidation, disconnection, and danger) correlated with genetic dispositions towards educational attainment, suggesting that the association could be in part due to selection effects, rather than a solely causal role of neighbourhood characteristics on educational outcomes (34).

We also leverage recent genetic discoveries to partition polygenic score effects and address two further, more specific, questions. First, we investigate how family environments mediate *cognitive* and *noncognitive* genetic effects on academic achievement over development. We decompose the polygenic signal in educational attainment into a cognitive and a noncognitive component (35). Our previous work (36) has shown that a PGS of noncognitive skills predicted variation in achievement across compulsory education and that its predictive power increased substantially over academic development, reaching effect sizes comparable to those observed for cognitive performance. As such, we investigate potential developmental effects in the environmental mediation of PGS effects on academic achievement separating cognitive and noncognitive genetics.

Second, we examine environmental mediation of polygenic score effects considering differences between siblings (37). Within-sibling analyses rely on how the transmission of alleles from parents to offspring is randomized during meiosis, such that siblings have an equal probability of inheriting any given allele, independently of environmental processes. Therefore, genetic differences between siblings are thought to be free from environmental influences shared by the siblings, which include passive gene-environment correlation. Differences at the within-sibling level thus are likely to reflect how each sibling perceives, evokes, and shapes the family environments (35). These additional analyses will allow us to dig deeper into the mechanisms through which family environments might contribute to the strengthening of the association between genetic propensity and academic outcomes over development (38,39). Evidence of mediation effects observed between families, but not between siblings, would be more consistent with passive gene-environment correlation processes, while environmental mediation observed when looking at differences between siblings would suggest evocative/active gene-environment correlation processes.

In summary, the current study answers three core research questions: First, do family environments mediate polygenic score effects on academic achievement over development? If so, which family environments matter and when in development? Second, do environmental mediation effects differ between cognitive and noncognitive genetics? Third, are environmental mediations of polygenic score effects observed also when considering differences between siblings? The protocol for the current study was preregistered with the Open Science Framework (OSF) and can be accessed at the following link: https://osf.io/tyf4v/. Deviations from the pre-registered protocol are described in **Supplementary Note 1.**

## Methods

### Participants

Our participants were twins enrolled in the Twins Early Development Study (TEDS) (40). TEDS has collected data from twins born in England and Wales between 1994 and 1996 and their parents at several points during childhood, adolescence, and early adulthood, starting from birth. Over 13,000 twin pairs took part in the first data collection, and nearly 30 years on, over 10,000 families remain active members of TEDS. The TEDS sample remains largely representative of the UK population for their generation in terms of ethnicity and socio- economic status (14). The subsample included in the current analyses consisted of 5151 individuals whose families had contributed data on academic achievement, family environment and who had genotype data available. The sample included 51% female and 49% male. We considered data collected over four waves: age 7 (Mean age = 7.15, age 9 (Mean age = 9.03), age 12 (Mean age = 11.53), and age 16 (Mean age = 16.31). Participants with severe medical, genetic, or neurodevelopmental conditions were excluded from our analyses. The sample size fluctuated between 5151 and 1439 due to incorporating distinct variables.

### Measures

Data were collected using questionnaires and tests administered to parents, teachers, and the twins themselves by post, telephone, and online, as described in detail in this overview of the TEDS study (41) and the TEDS data dictionary (https://www.teds.ac.uk/datadictionary/home.htm).

#### Academic achievement

Academic achievement was measured at ages 7, 9, 12 and 16 as a composite of academic performance across two subjects: English and mathematics. At ages 7, 9 and 12, data were provided by the teachers who assessed students’ performance based on the UK National Curriculum guidelines designed by the National Foundation for Educational Research (NFER; http://www.nfer.ac.uk/index.cfm) and the Qualifications and Curriculum Authority (QCA; http://www.qca.org.uk).

At age 16, academic achievement was measured as the mean grade score for the General Certificate of Secondary Education (GCSE) passes. GCSEs are standardized tests taken at the end of compulsory education, which in the UK is at age 16. The exams are graded on a scale ranging from A* to G, with a U grade assigned for unsuccessful attempts. The grades were coded on a scale from 11 (A*) to 4 (G, the lowest passing grade), and the mean of the grade obtained across the GCSE passed subjects was used as our measure of academic achievement at age 16. Data on GCSE performance were collected from parental and self-reports. Our previous research has shown that teacher ratings and self-reported GCSE grades are valid, reliable, and correlate very strongly with standardized exam scores taken at specific moments in the educational curriculum (Key Stages) obtained from the National Pupil Database (42).

#### Family environments

Data on family environments were collected from the twins and their parents at ages 7, 9, 12 and 16. In line with Bronfenbrenner’s ecological systems theory (43), we considered both wider contexts related to the family environment (i.e., neighbourhood characteristics) and proximal aspects of the family environment (e.g., socioeconomic status, parenting, home environment).

##### Neighbourhood characteristics

We obtained data on each family’s neighbourhood characteristics through geocoded data linkage with administrative data, which showed high consistency throughout development (see (44) and **Supplementary Note 2** for a detailed description). Postcodes were obtained from the TEDS families in 1998, 2008 and again in 2018. The 1998 and 2008 postcodes were for parent addresses, and as such were each linked to one set of data per family. The 2018 postcodes were for twin addresses, and the data are linked to individual twins. However, in most families both twins still had the same family address and postcode, and in these cases the linked data are identical for both twins. Administrative data included information on a broad range of intercorrelated neighbourhood characteristics (see **Supplementary** Figures 1**- 3**). The census data were downloaded from the Nomis web site, maintained by University of Durham, and operating on behalf of the *Office for National Statistics (ONS)*. Two sets of census variables were downloaded and linked: from the 2001 census and from the 2011 census. Census data included variables covering a wide range of characteristics of the resident population linked to each post code, such as the age and socio-demographic structure of the population, household spaces and types, occupancy, health (a full list of the census variables that were linked to the TEDS dataset can be found at the following link https://www.teds.ac.uk/datadictionary/pdfs/postcodes/postcode_linked_census.pdf). The pollution data were downloaded from *UK Air*, part of the UK Government’s *Defra* web site (Department for Environment Food & Rural Affairs): https://uk-air.defra.gov.uk/data/pcm-data. They provide measures of common gaseous and particulate atmospheric pollutants in given locations during a given year, including for example Nitrogen dioxide (NO2), Benzene and Ozone. A full list of the pollution variables linked to the TEDS dataset can be found at the following link: https://www.teds.ac.uk/datadictionary/studies/measures/postcode_linked_data.htm#pollution.

Given the large number of variables linked to each postcode, to reduce the dimensionality of the data, we conducted exploratory and confirmatory factor analyses (see Supplementary Figures 4-7), which resulted in the creation of six composites that measured features of the neighbourhood environment (see Supplementary Figures 4-7): (1) occupancy rating (indicating whether on average households in the neighbourhood had the required number of bedrooms, more (under-occupied) or less (overcrowded)); (2) health rating (indicating the proportion with general health rating (good, bad and fair) and proportion households with central heating), (3) household size (e.g., mean household size, mean population age, and proportion of households with lone parent plus children), (4) population in households (e.g., proportion of dwellings occupied, tenure proportion of social housing, and number of people living in the household), (5) qualification level (proportion with different levels of qualifications), and (6) pollution (e.g., annual mean particulate matter < 10). These six aspects of the neighbourhood environment have been previously shown to relate to children’s health and wellbeing (45,46), and educational and socioeconomic outcomes (47) to varying degrees.

##### Proximal home environments

***Positive and negative parental feelings*** were measured using 7 items derived from the Parent Feelings Questionnaire (48) which included questions on both positive and negative feelings a parent experiences about each child (e.g., positive item of “*Do you generally feel quite happy about your relationship with the ELDER twin?”* and negative item of “*Does the ELDER twin ever make you feel frustrated?*”). The 7 items were rated on a 4-point scale (in which 1 = *never* and 4 = *often*) for the firstborn twin. After parents answered questions about the firstborn twin, they were then asked: “do you feel this more or less often with the younger twin?” rated on a 3-point scale ranging from 1 = *more* to 3 = *less*. The same scales were collected at ages 7, 9 and 12. A composite score was created by summing the items (requiring at least 4) and reversing where necessary. At ages 9 and 12, each twin also reported their perception of parental feelings. The twins answered 7 questions (e.g., ‘*My Mum/Dad gets impatient with me*’ and ‘*My Mum/Dad finds me funny – I make him/her laugh*’) on the same 3-point Likert scale (0 = *often*, 1 = *sometimes*, 3 = *rarely or never*).

***Harsh parental discipline*** was assessed using the mean of four questionnaire items adapted from a semi-structured interview (49) asking parents about their discipline strategies when their child misbehaved. The questionnaire included negative discipline: shouting, sending the child to his or her room or withdrawing privileges, smacking, or restraining, and ignoring the child when he/she is misbehaving. The four items were rated on a 4-point scale (1 = *never* to 4 = *often*) for the firstborn twin. After parents answered questions about the firstborn twin, they were then asked, “Do you do this more or less often with the younger twin?”. This was rated on a 3-point scale ranging from 1 = *more* to 3 = *less*. Parents provided information on their discipline strategies when their children were 7, 9 and 12 years old. At ages 9 and 12, each twin also provided information on their parents’ discipline strategies by answering the same question as their parents.

***Chaos at home***. The degree of chaos at home was assessed by parents using a short version of the Confusion, Hubbub, and Order Scale (CHAOS) (50) which includes items that ask participants to rate the extent to which they live in a disorganized and noisy household.

Parents and children answered questions such as “*You can’t hear yourself think in our home*” and “*The atmosphere in our house is calm*” on a 3-point Likert scale (0 = Not true, 1 = Quite true, 2 = Very true). Parent reports were available at ages 9 and 12, and twin self-reports were available at ages 9, 12, and 16.

***Supportive home environment.*** At age 9, parents reported on several aspects of the home environment considered to stimulating for children’s development. A composite score was computed as the standardised mean of 3 parent-rated items: (1) *How many books in the home*, (2) *How often had the child been taken to the museum in the past year,* and (3) *There is a computer at home that is used by child*. Items were scored on a three-point Likert scale (0= not true, 1= somewhat true, 2= certainly true), with higher scores indicating a more stimulating home environment. The average correlation between the three items was 0.38.

Details of the creation of the *stimulating home environment scale*, including exploratory factor analysis and correlations between items, are included in **Supplementary Note 3** and **Supplementary** Figures 7-9.

***TV consumption.*** At age 9, a composite measure describing each child’s TV consumption was calculated as the standardised mean of self-rated items: (1) *On a normal school day how many hours of television does your child watch?* (2) *On a normal weekend day how many hours of television does your child watch?* Items were scored on a six-point Likert scale (0= 0 hours, 1= 1 hour up to 5 = 5 or more hours). Higher scores indicated more TV consumption in the home (Cronbach’s α= 0.67). Details of the TV consumption scale creation, including exploratory factor analysis and correlations between items, are included in **Supplementary Note 2** and **Supplementary** Figures 8-10.

***Parental monitoring and parental control*** were assessed using a set of six and eight items, respectively, reported by each of the twins at age 16. The questionnaires were both drawn from the NICHD Early Childcare and Youth Development Study (51). The twins rated the level of *parental control* in their family answering questions about who makes decisions about different activities, for example: “*Whether you can go out to meet friends*” and “*How you dress*”. The scale ranged from (1 = My parent(s) decide to 5 = I decide all by myself). The twins provided data on the level of *parental monitoring* by rating how much a parent or another adult in their home knew about different activities, including “*Who you spend time with*” and “*Where you go right after school*”. The scale ranged between (1 = Doesn’t know to 4 = Knows everything).

***Life events*** were measured through parent reports at age 9. A composite score was created by summing the number of significant life events experienced by each of the twins separately, with a higher score indicating a greater number of stressful life events. The scale consisted of 17 items asking parents to report on meaningful life experiences such as such as parents’ divorce or separation, death of a grandparent, unemployment, and financial difficulties.

***Family socioeconomic status (SES).*** Data on family SES were collected when the twins were 7 and 16 years old. At age 7, the family SES composite included data on parents’ occupational position (assessed by the Standard Occupational Classification 2000), educational qualifications, and maternal age at first birth. At age 16, family SES was calculated with a mean composite of standardized household income, maternal and paternal education level, and maternal and paternal occupation. Data on family income were also available when the twins were 9 years old.

##### Polygenic scores

After applying DNA quality control procedures recommended for chip-based genomic data (52), we constructed genome-wide polygenic scores (PGS) using summary statistics derived from five genome-wide association studies: educational attainment (16), cognitive ability (36,53) and noncognitive skills (35,36). Each PGS was calculated as the weighted sum of the individual’s genotype across all single nucleotide polymorphisms (SNPs). We used LDpred1 (54) to adjust for linkage disequilibrium. See (11) for a detailed description of our analytic strategy used to calculate PGS.

### Analytic strategies

#### Data preparation

All environmental measures were regressed on age and sex to control for their potential confounding influence. Polygenic scores were regressed on 10 principal components of ancestry and genotyping chip. The standardized residuals from these regressions were used in all analyses. Because some variables were skewed, analyses were repeated on square root transformed data (distributions and correlations between untransformed and transformed composites are presented in **Supplementary** Figures 11-13) and results were highly consistent.

#### Construction of latent factors measuring broader dimensions of the family environment

We applied exploratory factor analysis (EFA) (55) to examine the dimensionality of the family environment measures at different developmental stages. We performed EFA using *psych* for R (56) including the environmental measures available at each age. Based on the EFA results, we tested and created latent composites of correlated dimensions of environmental exposures using confirmatory factor analysis (CFA) (56) in *lavaan* for R (57). We examined model fit indices (**Supplementary Note 4**) to determine the goodness of fit of each model.

After exploratory and confirmatory factor analyses, dimensions of environmental exposure were constructed for all participants using CFA. Full Information Maximum Likelihood was used to account for data missingness. We constructed latent factors that captured broader dimensions of the family environment separately for each age and extracted factor scores that were used in subsequent analyses (see Results).

Specific procedures of EFA and CFA are illustrated in **Supplementary Note 4**, Correlation matrices between environmental variables at each age are presented in **Supplementary** Figures 14-17, scree plots are presented in **Supplementary** Figure 18 and factor structures yielded by each EFA are illustrated in **Supplementary** Figures 19-22. CFA models are illustrated in **Supplementary** Figure 23, and model fit indices are presented in **Supplementary Table 1**. All environmental composites and methods used for their construction are illustrated in **Supplementary** Figure 24. Correlations between the environmental composites are presented in **Supplementary** Figure 25.

#### Mediation analyses

After removing outliers (scores outside +/- 4 standard deviations), we conducted Structural Equation Modelling (SEM) for mediation analyses (58) using *lavaan* for R. A detailed description of the mediation models is presented in **Supplementary Note 5**. Mediation models allowed us to partition polygenic score effects on academic achievement into direct and indirect effects (i.e., effects mediated by exposure to each family environment).

We conducted mediation models considering the five polygenic scores and four academic achievement outcomes, selecting mediators for which data were collected at the same wave as the academic achievement outcome. For example, when considering academic achievement at age 7, we examined the effects mediated through supportive parenting and socioeconomic status, the two environmental measures collected when the children were 7 years old. We applied Benjamini-Hochberg false discovery rate correction (FDR) to account for multiple testing. In these analyses, we accounted for non-independence of observations in the sample (i.e., relatedness) by randomly selecting one twin out of each pair.

We applied *two-mediator mediation models* (59) to extend our investigation of mediation effects and examine whether the mediating role of family environment was simply driven by family SES. We repeated our mediation models considering each family environment jointly with family SES.

#### Multi-level mediation analysis: separating between from within-family effects

We separated within from between-family effects using *1-1-1 two-level mediation models* (60). This statistical model allowed us to examine the indirect effect of a predictor on an outcome by introducing mediation clustered data. For these analyses, we clustered our data by family, with each family corresponding to a cluster of two members (the two dizygotic twins). Applying 1-1-1 multilevel mediation models, we were able to separate between and within-sibling polygenic score effects while also separating mediation effects. Because the within-siblings PGS association is free from the effects of passive gene-environment correlation and demographic confounders, which are captured at the between-siblings level, this allowed us to test whether the mediating role of family environments was in line with the possibility of passive or evocative/active gene-environment correlation, or both.

Only dizygotic (DZ) twins were included in these analyses as we aimed to examine how within-siblings’ differences in polygenic scores predicted differences in academic achievement through differences in the family environment. Monozygotic (MZ) twin pairs could not be included as their polygenic scores do not differ within families. Furthermore, these analyses could only be performed for family environments that differed between the two twins, for example, reports of parenting and home chaos, but not for measures that were the same for both twins, such as family SES.

## Results

### Creating broader environmental measures

We applied exploratory and confirmatory factor analysis (see Methods) to derive broader composite measures of the family environment that could reflect the correlation between multiple aspects of the neighbourhood and home environment (see **Supplementary** Figures 1-10 and 14**-24; Supplementary Note 4).** From these analyses, we extracted the following higher-order dimensions of the family environment.

*Supportive parenting.* At age 7, we created a measure of supportive parenting, which was constructed as the mean composite of two parent-rated scales: (1) Positive parental feelings (reversed when necessary) and (2) a reverse-coded composite of the harsh parental discipline scale.

*Harsh parenting and chaos.* Based on our exploratory and confirmatory factor analysis results (**see Supplementary** Figures 20 and 21), we extracted a measure of “harsh parenting and home chaos”. This factor loaded three scales: (1) negative parental feelings, (2) harsh parental discipline and (3) chaos at home. We found a great deal of consistency in model fit across different informants and ages; therefore, we created this broad composite of *harsh parenting and chaos* for both parent and child-reported family environments at ages 9 and 12. We exported factor scores for these four dimensions (Parent and self-reported harsh parenting and chaos at age 9 and Parent and self-reported harsh parenting and chaos at age 12).

*Supportive home environment.* Based on our exploratory and confirmatory factor analysis results, at age 9, we also extracted a broader factor that we called “supportive home environment” on which loaded three parent-reported measures: (1) a composite score of the stimulating home environment scale, (2) household income, and (3) parental marital status (see **Supplementary** Figure 20).

*Parental monitoring and chaos* at age 16 were constructed as a mean composite of two self-reported scales of parental monitoring and chaos at home, which correlated moderately negatively (r = −0.23) (**Supplementary** Figure 22).

*Quality of the neighbourhood.* The results of exploratory and confirmatory factor analyses (see **Supplementary** Figures 5-7) led to the creation of six latent composites that measured broader aspects of the neighbourhood environment: (1) occupancy rating (2) health ratings, (3) household size, (4) population in households, (5) qualification level, and (6) pollution level.

These broader dimensions of the family environment were taken further into our main analyses. However, analyses were also conducted on each individual environmental measure (Supplementary Information). Descriptive statistics of all measures are presented in **Supplementary Table 2**.

### Family environments correlate with polygenic scores for educational attainment (EA), cognitive (Cog) and noncognitive (NonCog) skills

Consistent with previous work (35,36), we found that PGSs correlated with academic achievement across development and that associations became stronger over the course of compulsory education, particularly for the EA and NonCog polygenic scores. For example, the correlation between the EA PGS and academic achievement increased from 0.20 at age 7 to 0.36 at age 16 (Figure 1 and **Supplementary Table 5**). Correlations between individual family environmental measures and academic achievement are presented in **Supplementary Tables 6a-7d**.

When examining the association between PGSs and family environments, we observed the strongest positive associations with family socioeconomic status (SES), measured when the twins were 7 years old (e.g., EA, *r*= 0.31, *p* < = 0.001, 95% CI [0.28, 0.34]) and 16 years old (e.g., EA, *r* = 0.30, *p* < = 0.001, 95% CI [0.26, 0.35]; see **Figure 1** and **Supplementary Table 3a**).

**Figure 1:**
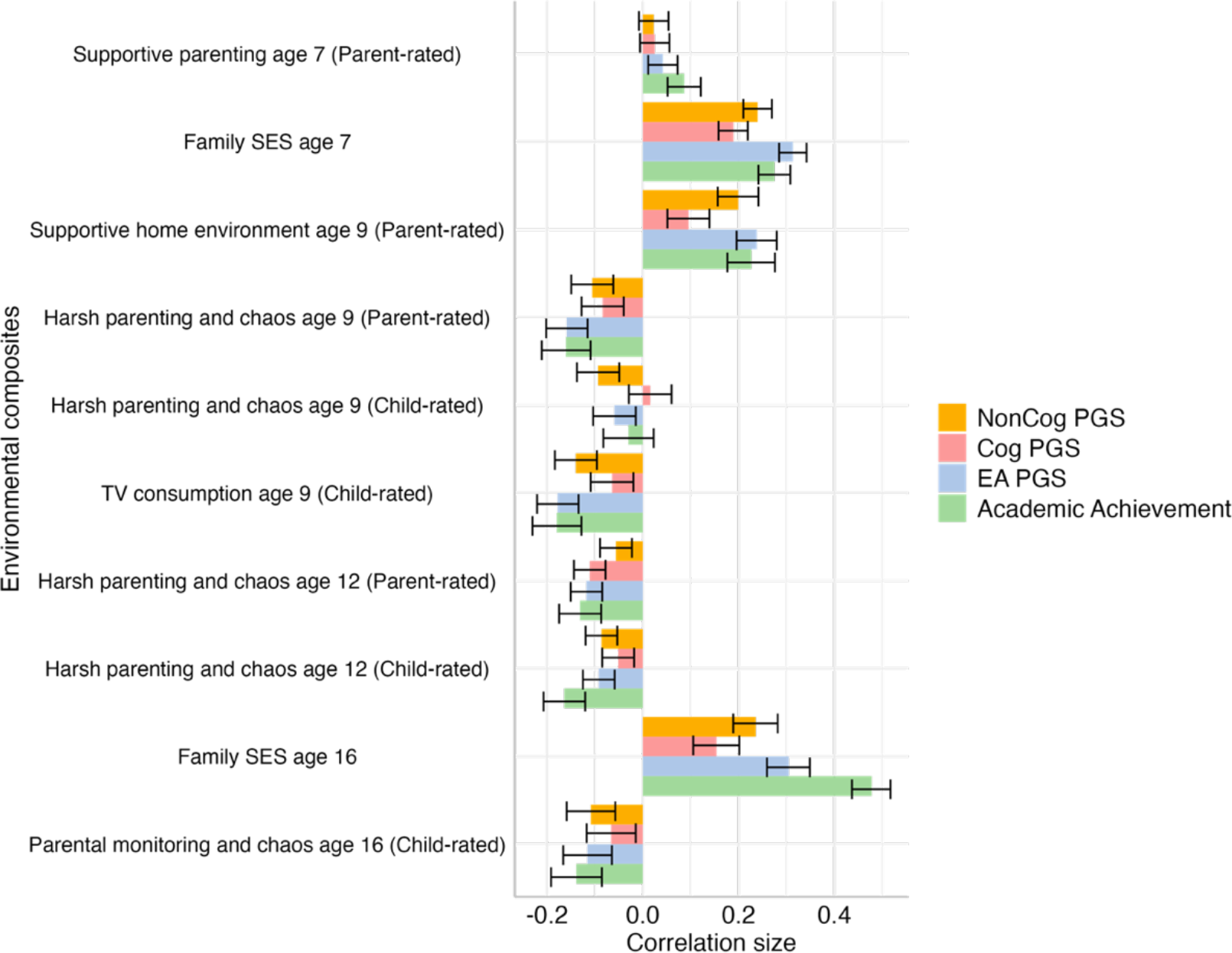
Correlations between environmental composite measures, children’s academic achievements and cognition and education-related polygenic scores. The measure of academic achievement included was contemporaneous to each environmental measure. EA PGS = educational attainment polygenic score. Cog PGS = Cognitive performance polygenic score. NonCog PGS = Noncognitive skills polygenic score. The length of each bar represents the size of the correlation coefficient and error bars indicate 95% confidence intervals.

Several other aspects of the family environment were also modestly correlated with all PGSs, for example, harsh parenting and chaos rated by parents (associations with EA were *r* = −0.16, *p* < = 0.001, 95% CI [-0.20, −0.12] at age 9 and *r* = −0.12, *p* < = 0.001, 95% CI [-0.15, −0.08]; at age 12,) and TV consumption at age 9 (*r* = −0.18, *p* < = 0.001, 95% CI [-0.22, −0.13] with the EA polygenic score). The supportive home environment composite at age 9 was also significantly associated with all PGSs (*r* = 0.24, *p* < = 0.001, 95% CI [0.20, 0.28] for EA, *r* = 0.10, *p* < = 0.001, 95% CI [0.05, 0.14] for Cog and *r* = 0.20, *p* < = 0.001, 95% CI [0.16, 0.24] for the NonCog polygenic scores). Correlation coefficients and p values for all environmental measures are reported in **Supplementary Tables 3a-3d**.

### Environmental measures correlate with measures of academic achievement across development

Environmental composites correlated with academic achievement across development with comparable effect sizes to those observed for the PGSs. For example, the correlation was 0.23, *p* < = 0.001, 95% CI [0.18, 0.28] between a supportive home environment at age 9 and academic achievement at the same age, and 0.28, *p* < = 0.001, 95% CI [0.24, 0.31] between family SES at age 7 and academic achievement at the same age (**Figure 1** and Supplementary Tables 4a-4d).

### Family environments mediate PGS effects on academic achievement across development

Given the associations observed between PGSs, family environments, and academic achievement, we conducted mediation models to examine the extent to which these aspects of the family environment mediated the prediction from genetic disposition to variation in academic achievement over development. We started by examining the role of more distal neighbourhood characteristics and continued to explore the role of aspects of the home environment more proximal to each child.

#### Quality of the neighbourhood

We first examined the role of neighbourhood characteristics (occupancy rating, health ratings, household size, population in households, qualification level, and pollution). We found significant and consistent, yet weak, mediation effects for selected neighbourhood measures. Neighbourhood occupancy, health, and household size mediated the prediction from the educational attainment polygenic score to academic achievement at ages 7, 12 and 16, but the average indirect effect was weak (average beta coefficient ß = 0.01 [95% CI, 0.01- 0.02]; see **Figure 2** and **Supplementary Table 8a).** Similar findings were observed when Cog and NonCog PGSs examined separately **(Supplementary** Figure 26 **and Supplementary Tables 8b-8c)**.

**Figure 2:**
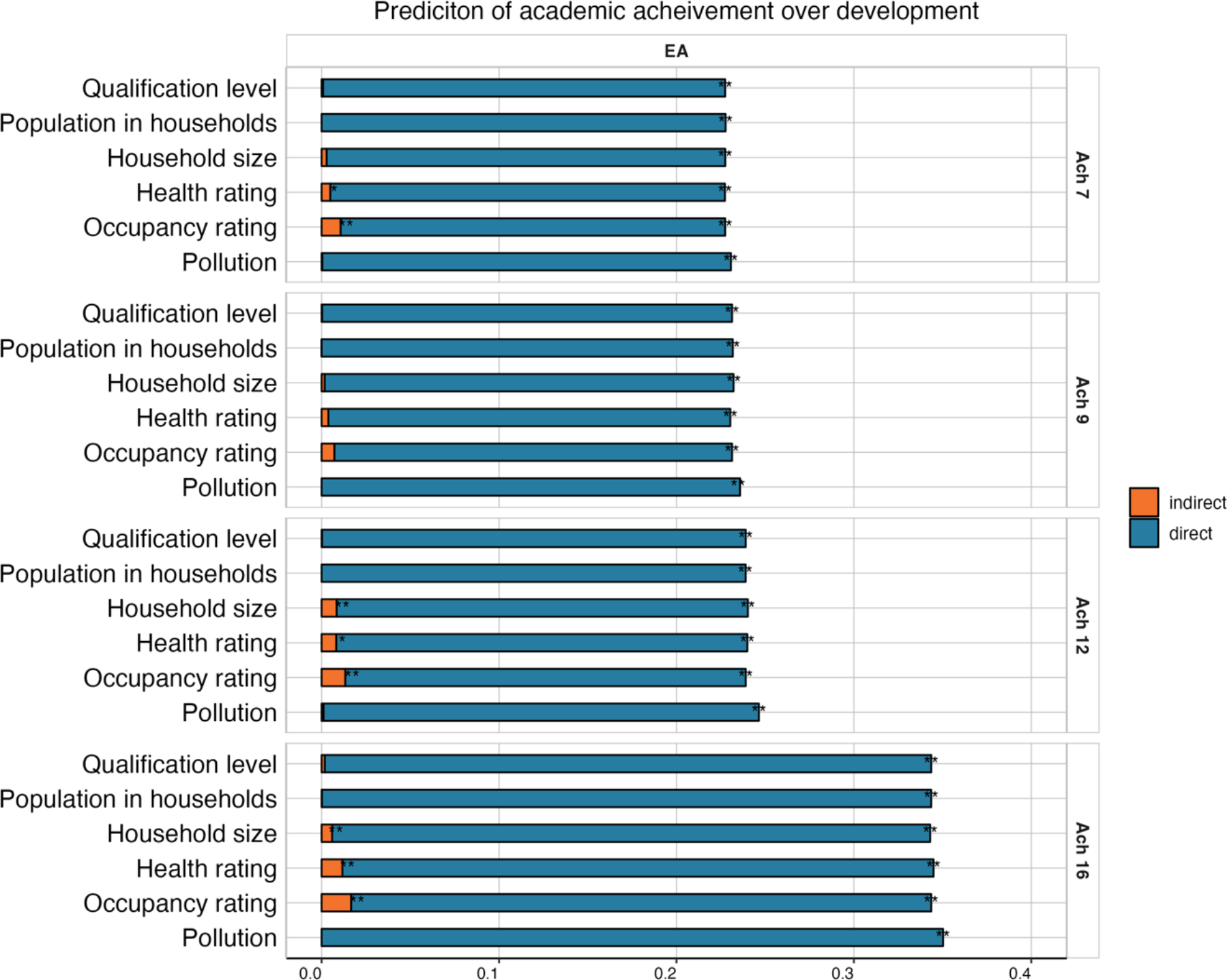
Mediating role of neighbourhood characteristics on the educational attainment polygenic score (PGS) prediction of academic achievement from age 7 to 16. The total length of each bar represents the prediction (standardised beta coefficient) from the educational attainment (EA) (16) PGS to academic achievement at ages 7 (top panel), 9, 12, and 16 (bottom panel). The blue portion of each bar shows the direct effect (i.e., not mediated by each neighbourhood measure), while the orange portion of each bar shows the indirect effect (i.e., mediated by each neighbourhood measure). * = p < 0.05, ** = p < 0.01 after applying FDR correction.

#### Home environments

We next examined whether more proximal aspects of the family environment could account for part of the genetic effects on academic achievement over development. We examined the role of family environmental contexts at multiple levels of granularity, moving from broad constructs that reflected commonalities across environmental measures to specific indices of the environmental contexts (61).

Figure 3 presents the results of mediation analyses for broader measures of the family context, including SES, supportive home environment and harsh parenting and chaos. When considering the pathway from the EA PGS to academic achievement over development, we found significant mediating effects for most environmental contexts, except for child-rated harsh parenting and chaos at age 9. The strongest indirect effects were found for SES at age 7 (ß = 0.07 [95% CI, 0.06-0.08]) and age 16 (ß =0.11 [95% CI, 0.09-0.13]), when SES mediated nearly 1/3 of the EA PGS prediction. A supportive home environment at age 9 (ß =0.05 [95% CI, 0.03-0.06]) was also found to have a substantial mediating role (∼ ¼ of the total prediction). Model estimates are presented in **Supplementary Table 9**.

**Figure 3:**
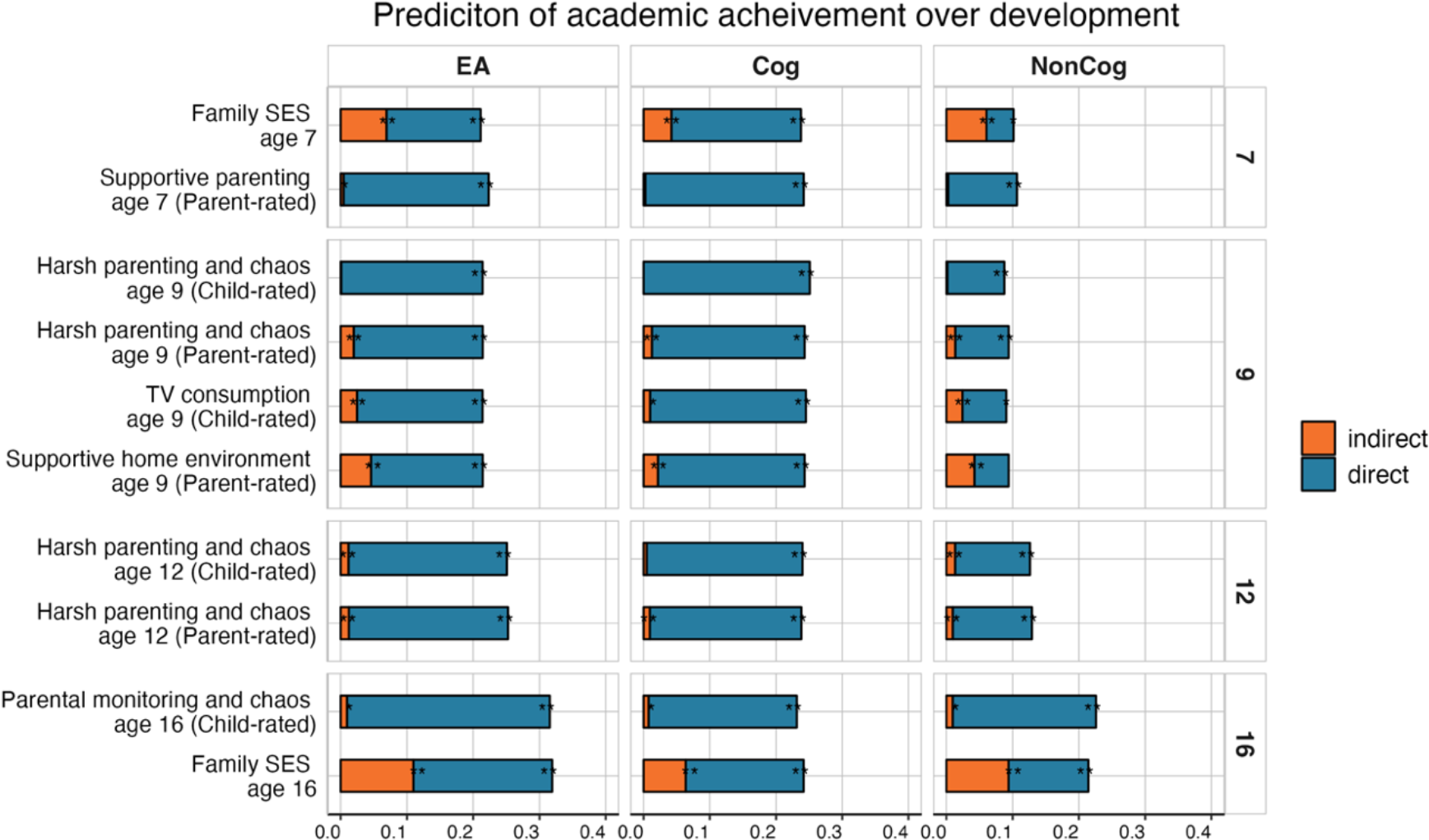
Polygenic scores (PGS) effects on academic achievement across compulsory education mediated by home environments. The total length of each bar represents the prediction (standardised beta coefficient) from the educational attainment (EA; (16)), cognitive (Cog; (53)) and noncognitive (NonCog; (35,36)) PGS to academic achievement at ages 7 (top panel), 9, 12, and 16 (bottom panel). The blue portion of each bar shows the direct effect, while the orange portion of each bar shows the indirect (i.e., mediated) effect. * = p < 0.05, ** = p < 0.01 after applying FDR correction.

Since we found significant mediating effects for the EA PGS, we examined whether these could be captured by cognitive or noncognitive PGSs. Figure 2 therefore shows the mediating effects of the family environment in the prediction from Cog and NonCog PGS to academic achievement over development. Although a similar pattern of results emerged for both Cog and NonCog PGSs, effects were stronger for the NonCog PGS prediction (e.g., the indirect effect of family SES was ß = 0.06 [95% CI, 0.04-0.09] for Cog and ß = 0.09 [95% CI, 0.07- 0.12] for NonCog; **Supplementary Tables 10a and 10b)**, particularly when considering the total PGS effect. Although PGS predictions were weaker for NonCog PGS, mediating effects approached or even exceeded half of the total PGS effect (e.g., for family SES at age 7; Figure 2 right panel).

Mediating effects for specific indices of the family environmental contexts were generally weaker, although many environments significantly contributed to the PGS effects on academic achievement at all ages. For example, home chaos across all measurements accounted, on average, for 11% of the total EA PGS effects (see details in **Supplementary** Figure 27 and **Supplementary Table 11**). A similar pattern of results emerged when we repeated the analyses with three other PGSs (for IQ (53), Cognitive and Noncognitive skills (40)). Results are presented in **Supplementary** Figures 28 and 29; **Supplementary Tables 12 and 13.**

### Controlling for the effects of SES using a two-mediator mediation model

Considering that SES was the strongest mediator of the PGS prediction of academic achievement at several developmental stages, and considering its correlations with several other aspects of the family environment, we tested whether our results were driven by family SES. To this end, we extended our mediation models to include family SES as an additional mediator and run two-mediator mediation models (see Methods). These models allowed us to test whether all other aspects of the family environment remained significant mediators after accounting for the role of family SES. Because family SES was measured at ages 7 and 16, for all models predicting achievement at ages 7, 9 and 12, we included family SES measured at age 7, while for the models predicting achievement at 16, we included a measure of family SES collected when the twins were 16 years old. Although we found that family SES played a significant role in mediating the PGS predictions of academic achievement, the indirect effects of other environmental measures (e.g., harsh parenting and CHAOS and supportive home environment) remained significant, albeit attenuated (**Supplementary** Figure 30 and **Supplementary Table 13**). Similar results were observed across all PGSs and at all developmental stages. (**Supplementary** Figures 31 and 32; **Supplementary Tables 14 and 15**).

### Separating mediation effects into between- and within-families to further investigate gene- environment correlation

Given the outcomes of our mediation analyses, which point to widespread gene-environment correlation in academic development, we applied multilevel mediation models (see Methods) to investigate whether family environments mediated the PGS-achievement relationship not only between but also within families. These analyses were only possible for those environmental measures that differed between siblings. The number of families included in the analyses ranged between 1278 to 3356, depending on the age of data collection and measures included. Specifically, family-level intraclass correlations (ICCs) ranged between 0.32 and 0.99 (**Supplementary Table 16**). As expected, PGS effects were attenuated at the within-family level (62). We also observed that nearly all mediation effects were captured at the between-family level for the EA PGS prediction of achievement across development (Fig 4 and **Supplementary Table 17**) and the same was observed for Cog and NonCog PGS effects (**Supplementary** Figures 33 and 34; **Supplementary Tables 18 and 19**).

**Figure 4:**
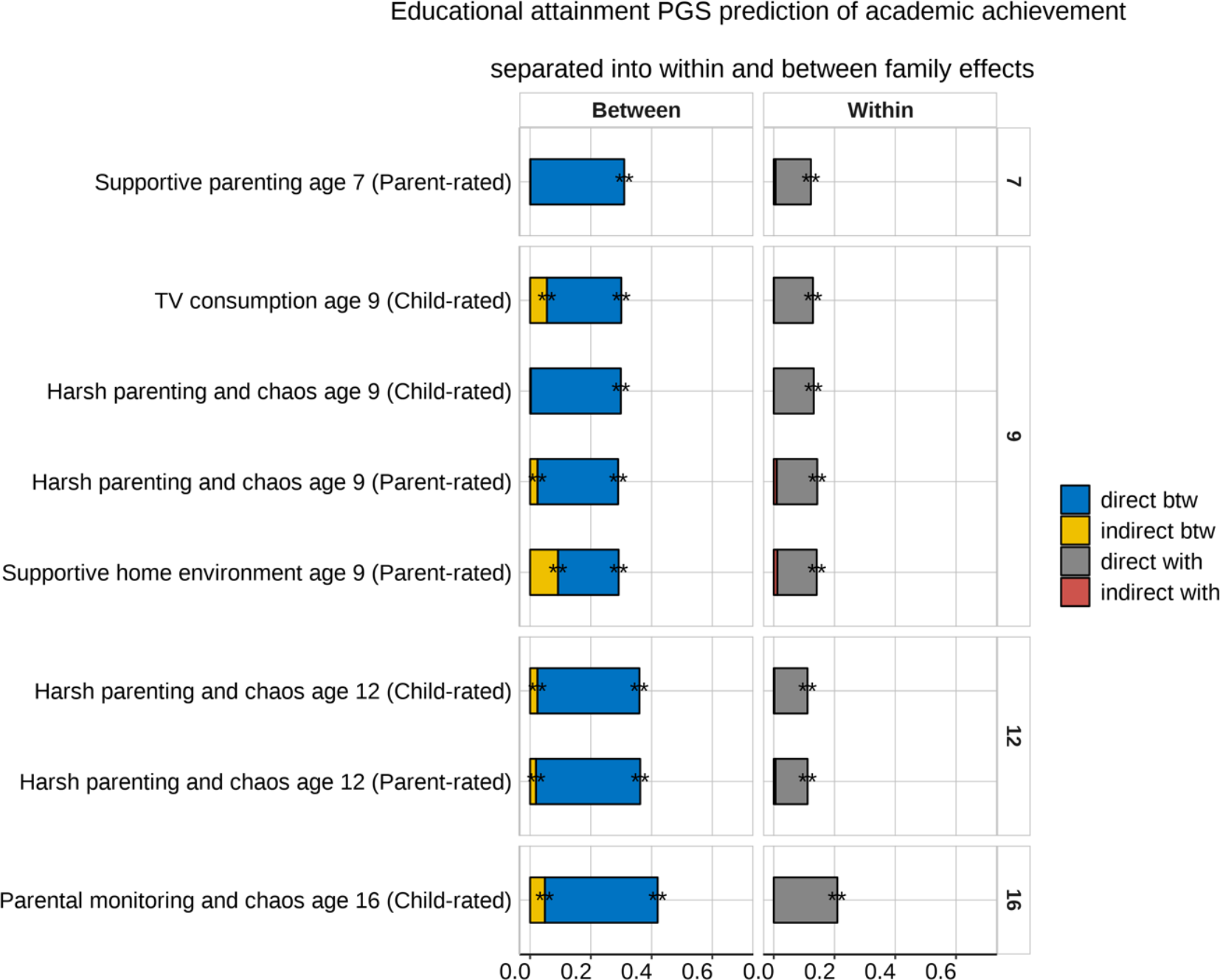
Educational attainment (EA) polygenic score (PGS) effects on academic achievement separated into between- and within-family effects mediated by family environments. The total length of each bar represents the effect of the EA PGS prediction (standardised ß coefficient) of academic achievement at ages 7, 9, 12 and 16, partitioned into between and within family effects. The blue and grey portions of each bar show the direct effects for the between and within-family levels, respectively. The yellow and red portions show indirect (mediated) effects for the between and within-family levels, respectively. * = p < 0.05, ** = p < 0.01 (FDR corrected).

## Discussion

The current study provides a systematic investigation of the role that family environments play in translating genetic disposition into observed individual differences in academic achievement over compulsory education. Three core findings emerged. First, we found evidence for widespread gene-environment correlation. Second, we found that family environments are more robustly linked to noncognitive genetic effects on academic achievement than cognitive polygenic score effects. Third, we found that the mediating role of family environments was nearly exclusively observed for between-family polygenic score effects, which is consistent with the hypothesis that family environmental contexts contribute to academic development via genetic nurture, rooted in passive gene-environment correlation processes. Passive gene-environment correlation suggests that parents shape environments for their children partly based on their genetic disposition. Our findings suggest that parents shape educationally relevant environments for their children, particularly in line with their disposition towards education-related noncognitive skills. In addition, parents might be more likely to shape these environments based on their genetic propensity rather than responding to every child’s specific genetic dispositions.

Our first set of results also provides finer-grained details on which environments matter for academic development. We found that, if compared to the effects of proximal family contexts, more distal aspects of the family environment, such as neighbourhood socio- economic status, health, and pollution, play a smaller role in translating genetic disposition towards education into observed individual differences in academic development. This is in line with previous work showing that selection effects rather than causal pathways are likely to underlie the association between neighbourhood deprivation and educational outcomes (34).

The effects of more proximal family contexts were consistent over development, both when considering broader composite scores, such as stimulating home environments, and when considering more fine-grained indices, for example, TV consumption. Family socio- economic status played the biggest role in mediating genetic effects on academic achievement, at all developmental stages. This is in line with previous research that has emphasised the importance of socio-economic factors in education, beyond genetics and cognitive ability (63). Our findings point to the importance of considering the complexity of how family environments, including socio-economic status, might contribute to academic development, complex processes that are not independent of, but in fact, correlated with genetic effects.

The current work also shows that, beyond socio-economic status, other aspects of the home environment, particularly parenting, contribute to academic development at all ages. This suggests that a supportive and stimulating family environment contributes to narrowing the achievement gap between children across socioeconomic brackets and levels of genetic disposition towards academic achievement (30,64). However, after accounting for socio- economic status, effect sizes were greatly reduced. Considering the substantial stability of socioeconomic factors and the stability of their effect on educational attainment across multiple generations (65), this is consistent with recent evidence suggesting that a part of the indirect genetic effects in education reflects dynastic effects that are common to families across more than one generation rather than characteristics specific to each nuclear family (66).

Our second main set of results highlighted how several environmental contexts exerted a greater mediating role when considering children’s genetic disposition towards noncognitive skills if compared to cognitive genetic dispositions. This is consistent with previous research finding significant genetic associations between noncognitive measures and academic achievement beyond cognitive skills (67,68). Our findings are also in line with research pointing to a greater role of family environments in contributing to genetic effects on noncognitive traits, such as personality and emotional stability, if compared to cognitive abilities (69). This indicates that parents not only create educational environments for their children that align with their genetic disposition towards cognitive abilities, but also shape these environments in line with their genetic dispositions towards socio-emotional skills.

With a third set of analyses, we aimed to delve deeper into gene-environment correlation mechanisms by separating between-family from within-family polygenic score effects. Genetic differences between siblings are likely to be free from environmental influences shared by them, which include passive gene-environment correlation. Consequently, significant mediation effects at the level of the within-family polygenic score prediction would index evocative (or active) gene-environment correlation that is driven by an individual child within a family. However, our results showed that most mediation effects were observed only between families, therefore consistent with passive gene-environment correlation processes. This is in line with previous research that found evidence for the role of parental investment in children’s educational outcomes operating via genetic nurture, which refers to how parents shape the family environment for their children partly depending on their own genetic dispositions (26,70). This is also consistent with previous work showing that genetic nurture effects on children’s academic and developmental outcomes are substantially mediated by prenatal maternal health and financial stability (71). Our results corroborate these findings by triangulating evidence using a different methodology. It is possible that, while we found that the family environment operates largely through passive gene-environment correlation processes, other environmental influences on academic achievement (e.g., school environment, socio-emotional factors, and close relationships) might operate through evocative and/or active processes, unique to each child rather than shared between families (72).

Several limitations should be acknowledged. First, the environmental variables used in our models are, at best, an imprecise representation of the actual family environments relevant to academic achievement. Similarly, polygenic scores are an imperfect and partial stand-in for additive genetic effects on academic achievement. Consequently, the results obtained from our mediation analyses might be confounded (73,74). Second, and related, it is possible that, in mediation analyses of polygenic score effects, the mediation pathway may be under-corrected for genetic confounding in the environmental variable, which could result in the genetic effects mediated by environmental risk factors to be overestimated (75).

Third, the current study was conducted in a UK-based sample, and it is unclear whether our findings would generalise to other populations characterised by different socio-contextual milieus. Fourth, the focus on White-European ancestry limits generalizability, however, recent multi-ancestry genome-wide association studies (76), and novel methods (77) are expanding the scope of genetic research to diverse populations, which can be used to address such gaps in future research. Fifth, although we examined the role that environmental factors play in academic achievement at several points during compulsory education, our mediators were cross-sectional which may not capture the evolving interplay between genetic and environmental factors over development. Longitudinal studies offer a richer perspective, tracing how early experiences might shape subsequent ones, a nuance potentially missed in our approach. We plan to extend our work and consider the cascading role of environmental influences longitudinally. Sixth, we investigated environmental mediation effects across the entire distribution of genetic disposition and academic achievement, but effects could be stronger when considering students at particularly high or low risk of underachievement.

Lastly, as new methodologies to partition between-family from within-family genetic effects emerge (78), we aim to extend our work and continue triangulating evidence across multiple methods.

To conclude, we provide evidence for the important role that aspects of the family environment, such as supportive parenting and a stimulating home environment play at every stage of academic development, beyond socio-economic factors. Our results suggest that parents shape environments that foster their children’s academic development largely based on their own genetic disposition, particularly towards noncognitive skills, through gene-environment correlation processes. These complex processes should be considered and controlled for when researching causes and correlates of individual differences in child development and learning.

## Code availability statement

The code is available at https://github.com/CoDEresearchlab/Achievement_Env_Mediation.

## Acknowledgements

This study was funded by a starting grant awarded to MM by the School of Biological and Behavioural Sciences at Queen Mary, University of London. QZ is funded by the QMUL- Chinese Scholarship Council joint PhD Scholarship. AG is supported by a Queen Mary School of Biological and Behavioural Sciences PhD Fellowship awarded to MM. TEDS is supported by a programme grant to RP from the UK Medical Research Council (MR/V012878/1 and previously MR/M021475/1), with additional support from the US National Institutes of Health (AG046938).

## Author contributions

These authors are contributed equally: QZ and AG.

Conceived and designed the study: M.M., Q.Z., A.G.. Analysed the data: Q.Z., A.G., M.M.. Wrote the paper: M.M., Q.Z, A.G., with helpful contributions from A.G.A., R.C.J.W., J.M.,

R.P. and K.R.. All authors contributed to the interpretation of data, provided feedback on drafts, and approved the final draft.

