## Supplementary Notes and Figures for "Gene-environment correlation: The role of family environment in academic development"

Note 1. Deviation from the preregistered analyses.
Note 2. Further information on neighborhood measures.
Note 3. Creation of Stimulating home environment and TV consumption scales.
Note 4. Exploratory and confirmatory factor analysis.
Note 5. Description of mediation models.

**Supplementary Figures**

Figure 1. Correlations between environmental pollution measures.
Figure 2. Correlations between neighbourhood quality measures.
Figure 3. Correlations between neighbourhood economy measures.
Figure 4. Scree plots of environmental pollution, neighbourhood quality and neighbourhood economy measures.
Figure 5. Factor structure of environmental pollution measures.
Figure 6. Factor structure of neighbourhood quality measures.
Figure 7. Factor structure of neighbourhood economy measures.
Figure 8. Correlations between parent and self-rated home environment items at age 9.
Figure 9. Scree plots of parent and self-rated home environment items at age 9.
Figure 10. Factor structure of parent and self-rated home environment items at age 9.
Figure 11. Distributions of cross-sectional composites.
Figure 12. Distributions of square root transformed cross-sectional composites.
Figure 13. Correlations between untransformed and square root transformed cross-sectional composites.
Figure 14. Correlations between parent-rated environmental variables at age 7.
Figure 15. Correlations between parent and self-rated environmental variables at age 9.

Figure 16. Correlations between parent and self-rated environmental variables at age 12.
Figure 17. Correlations between self-rated environmental variables at age 16.
Figure 18. Scree plots of parent and self-rated environmental variables at first contact and ages 7, 9, 12 and 16.
Figure 19. Factor structure of parent-rated environmental variables at age 7.
Figure 20. Factor structure of parent and self-rated environmental variables at age 9.
Figure 21. Factor structure of parent and self-rated environmental variables at age 12.
Figure 22. Factor structure of self-rated environmental variables at age 16.
Figure 23. CFA models of parent and self-rated latent cross-sectional composites at ages 9 and 12.
Figure 24. Cross-sectional composites.
Figure 25. Correlations between composites.
Figure 26. Cognitive and noncognitive PGS effects on academic achievement over development mediated by neighbourhood environments.
Figure 27. Educational attainment, cognitive and noncognitive PGS effects on academic achievement over development mediated by individual measures of the family environment.
Figure 28. Environmental mediation models using other cognitive and noncognitive polygenic scores. Cognitive and noncognitive PGS effects on academic achievement over development mediated by family environmental composites.
Figure 29. Environmental mediation models using other cognitive and noncognitive polygenic scores. Cognitive and noncognitive PGS effects on academic achievement over development mediated by individual measures of the family environment.
Figure 30. Indirect (environmentally mediated) *educational attainment* PGS effects on academic achievement before (pink) and after (blue) accounting for SES using two-mediators models.
Figure 31. Indirect (environmentally mediated) *cognitive* PGS effects on academic achievement before (pink) and after (blue) accounting for SES using two-mediators models.
Figure 32. Indirect (environmentally mediated) *noncognitive* PGS effects on academic achievement before (pink) and after (blue) accounting for SES using two-mediators model.
Figure 33. Environmentally mediated *cognitive* PGS effects on academic achievement across development, separated into within and between family effects.
Figure 34. Environmentally mediated *noncognitive* PGS effects on academic achievement across development, separated into within and between family effects.

#### Supplementary Note 1. Deviation from the preregistered analyses.

This study includes all the developmental, cross-sectional analyses preregistered at the following link: https://osf.io/tyf4v/. We are currently working on a follow-up study that extends our analyses to consider longitudinal mediators, therefore modelling stability and change in the environmental mediators.

#### Supplementary Note 2. Background information of neighborhood measures.

Of the 10469 families with recorded postcodes in both 1998 and 2005, 55% maintained the same address; in addition, 49% of 9803 families retained the same postcode from 1998 to 2010. Between 2005 and 2010, 86% of 10121 families had unchanged postcodes, underscoring the relative residential stability of the cohort. More details can be found in the TEDS data dictionary:
https://www.teds.ac.uk/datadictionary/studies/measures/postcode_linked_data.htm

#### Supplementary Note 3. Creation of the *stimulating home environment* and *TV consumption* scales.

To create the scales capturing variation in the home environment at age 9, we first explored the factor structure of parent-reported and self-reported data, separately. Examples of items related to the home environment included: '*The TV is on when the child is doing homework*', '*I discuss school activities with the child', 'My child reads for fun',* and '*Hours of TV watched per weekend day*'. Items were scored on a six-point Likert scale (0= 0 hours, 1= 1 hour up to 5 = 5 or more hours) with higher scores indicating more hours spent watching TV or doing enriching activities such as reading and going to museums. Correlations between parent and self-reported home environment items are illustrated in **Supplementary Figure 33**.

To explore the factor structure of the home environment items at age 9, we conducted an exploratory factor analysis (EFA) of parent and self-rated items. EFA analyses were conducted in *psych* for R (1, 2) and involved a sample of 3238 and 2941 independent twins, for parent and self-rated data respectively, created by randomly selecting one twin per pair. Results of the EFA are presented in **Supplementary Figure 34** and **Supplementary Figure 35**.

We adopted the data-driven approach and created parent-rated *Stimulating home environment* and self-rated *TV consumption* scales. The parent-rated *Stimulating home environment* scale was constructed as the standardised mean of parent-rated items: 1) *How many books at home, 2) Child was taken to museums in the past year* and 3) *Computer at home used by the child.* The self-rated *TV consumption* scale was calculated as the standardised mean of self-rated items: 1) *Hours of TV watched on a school day* and 2) *Hours of TV watched on a weekend day*, as guided by the EFA results. Cronbach's α= 0.38 for the parent-rated measure and 0.67 for the self-rated scale.

**Supplementary Note 4. Exploratory and confirmatory factor analysis.**

We examined model fit indices, including the Comparative Fit Index, Tucker-Lewis Index, Akaike Information Criterion, Bayesian Information Criterion and Root Mean Square Error of Approximation to determine the goodness of fit of each model.

EFA analyses were conducted using a sample of up to 11,497 randomly selected unrelated twins and CFA models were tested on the other half of the randomly selected sample (the other randomly selected sibling). Correlation matrices between environmental variables at each age are presented in Supplementary Figures 14-17, scree plots are presented in Supplementary Figure 18 and factor structures yielded by each EFA are illustrated in Supplementary Figures 19-22.

CFA models are illustrated in Supplementary Figure 23 and model fit indices are presented in Supplementary Table 1. Because successful convergence of CFA models requires > 2 manifest variables to define the latent composite, standardized mean scores were derived for latent constructs comprising < 3 manifest variables and for those that showed convergence problems or poor model fit. All cross-sectional composites, along with the methods used for their construction, are illustrated in Supplementary Figure 24. Correlations between the cross-sectional composites are presented in Supplementary Figure 25.

**Supplementary Note 5. Description of SEM-based mediation models.**

The formula for this mediation model for subject $i$ ($1 \leqslant i \leqslant n$) is given by:

$$z_i = \beta_0 + \beta_{xz} x_i + \varepsilon_{zi},$$
$$y_i = \gamma_0 + \gamma_{zy} z_i + \gamma_{xy} x_i + \varepsilon_{yi}$$

It is posited that the error terms ($\epsilon z_i$, $\epsilon y_i$) are uncorrelated, a critical assumption for causal inference when conducting mediation analysis. The assumption of multivariate normality for the error terms is also made, as it is an essential precondition for defining direct, indirect, and total effects. It should be highlighted that the two structural equations are interconnected, and the inference drawn from them is concurrent, rather than from two separate standard regression equations. More information can be found here (3).

**Population-level mediation analyses**

We conducted mediation analyses (Baron & Kenny, 1986; Preacher & Kelley, 2011) using the lavaan package for R to examine the direct and indirect effects of the prediction from genetic disposition (quantified as the educational attainment, cognitive and noncognitive skills polygenic scores) to manifested variation in academic achievement (Figure 1).

Mediation models estimate the indirect effect of a predictor (X) on the outcome (Y) via a mediator (M; i.e., an intervening variable –in this project family environments) by regressing M on X and regressing Y on both X and M using two separate equations:

$$1)\ M_i = d_{M.X} + aX_i + e_{M.Xi}$$

Where M$i$ is the mediator for individual $i$; $d_{M.X}$ is the intercept for the mediator (M); $a$X$i$ is the slope of M regressed on the predictor (X) and $e_{M.Xi}$ is the measurement error for individual $i$.

$$2)\ Y_i = d_{Y.MX} + bM_i + c'X_i + e_{Y.MXi}$$

Where Y$i$ is the outcome for individual $i$; $d_{Y.MX}$ is the intercept for the outcome (Y); $bM_i$ is the slope of the outcome (Y) regressed on the mediator (M) controlling for the predictor (X); c'X$i$ is the slope of the outcome (Y) regressed on the predictor (X) controlling for the mediator (M) and $e_{Y.MXi}$ is measurement error for individual $i$.

The indirect effect of the predictor on the outcome (i.e., the mediation effect) is defined by a^ × b^, with the sample estimate signified by the circumflex ("^").
When $a^\wedge \times b^\wedge = c^\wedge - c^\wedge{}'$, then $c^\wedge = a^\wedge \times b^\wedge + c^{\wedge\prime}$. Implementing SEM allows for, a^ and b^ can be derived simultaneously and for testing more complex models with latent class predictor, outcomes, and mediators.

We applied a sandwich correction to account for non-independence of observation (i.e. relatedness).

### Supplementary Figures

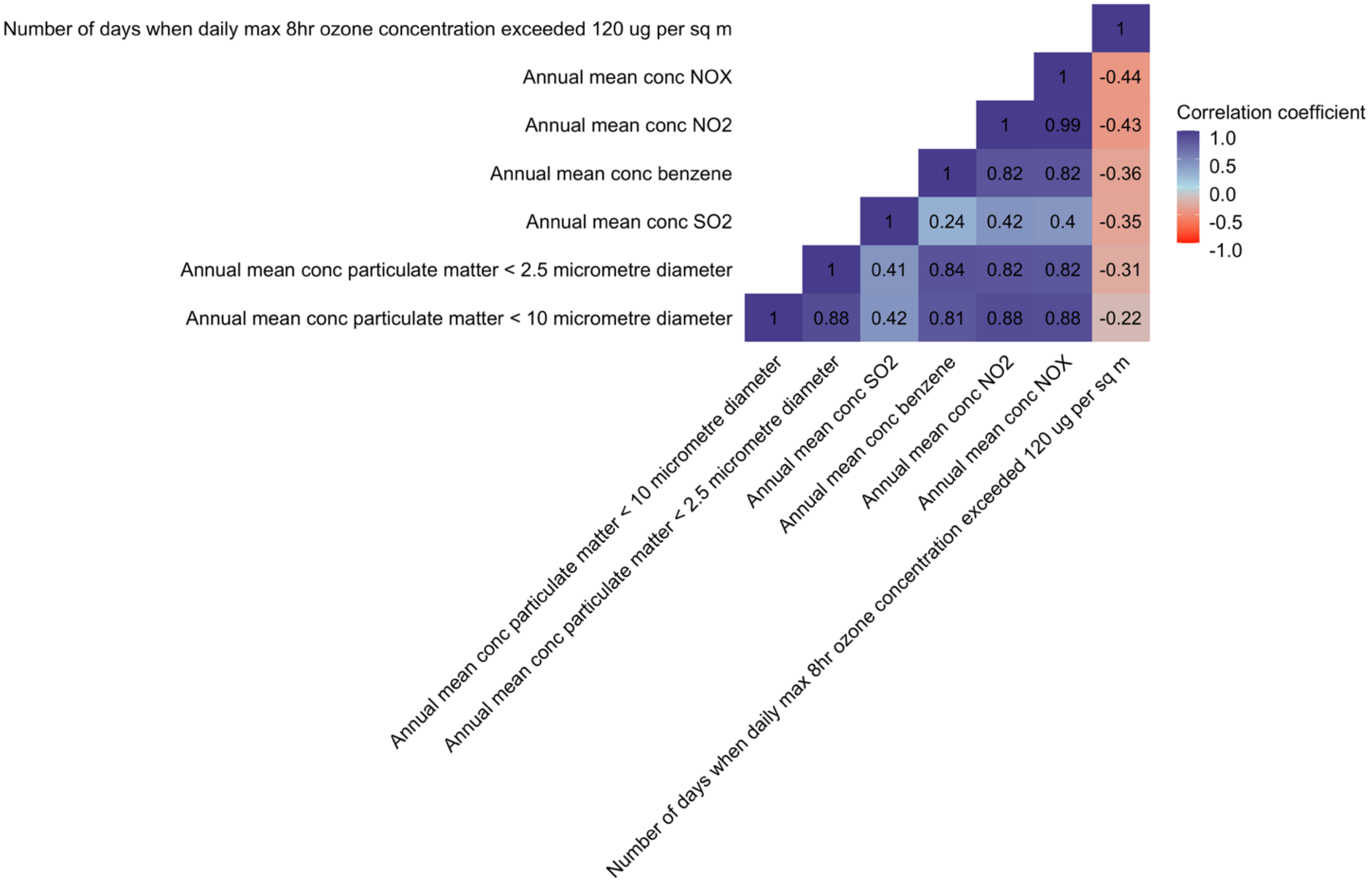

**Supplementary Figure 26.** Correlations between environmental pollution measures.

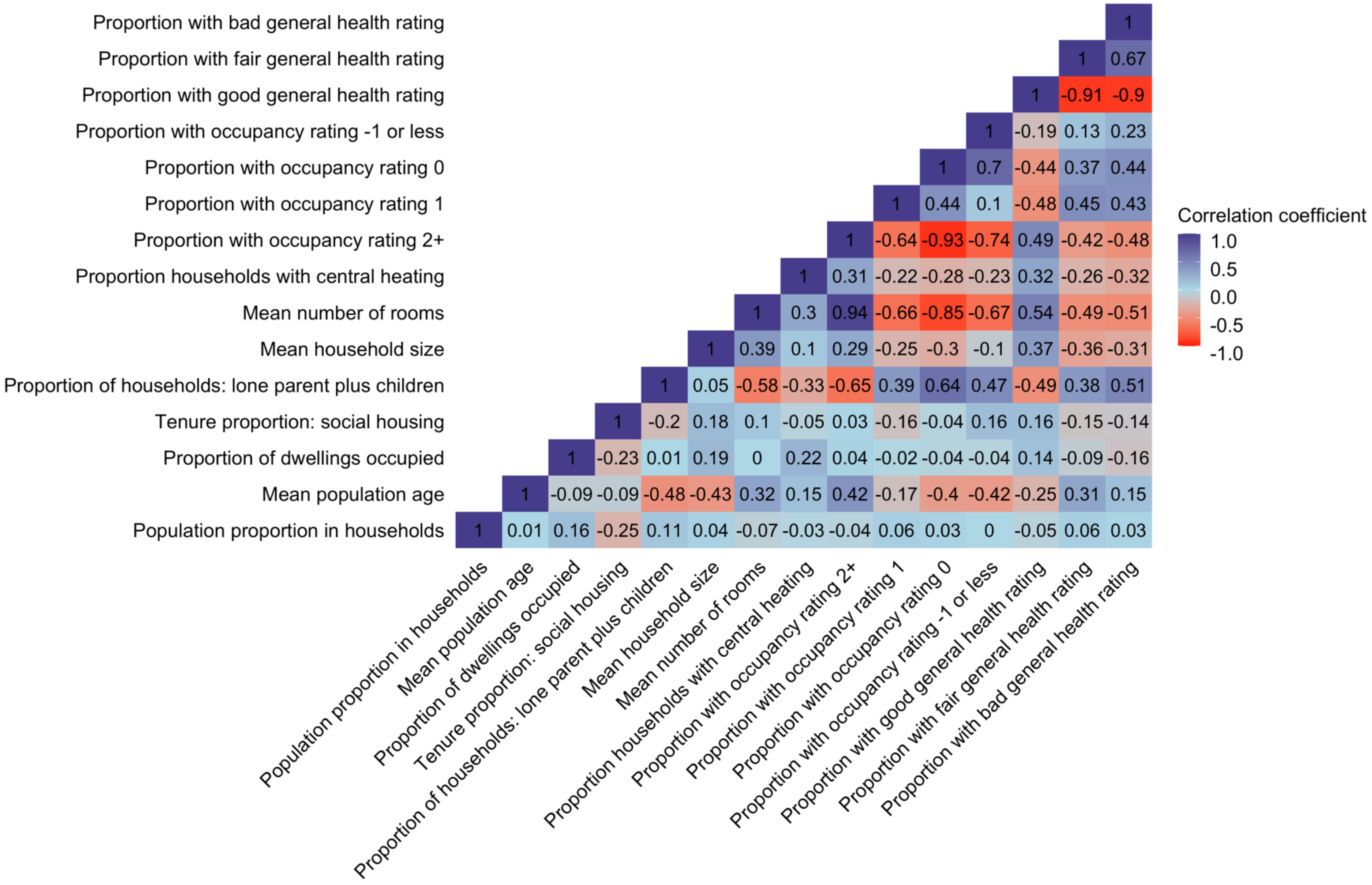

**Supplementary Figure 27.** Correlations between neighbourhood quality measures.

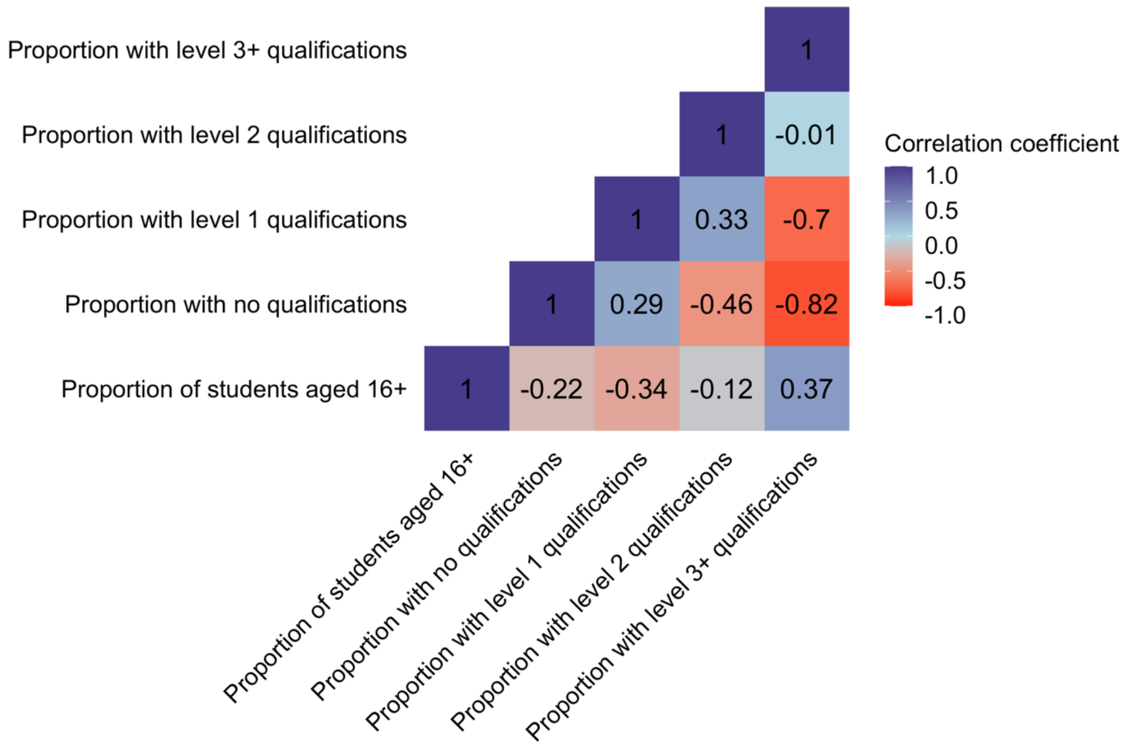


**Supplementary Figure 28.** Correlations between neighbourhood economy measures.

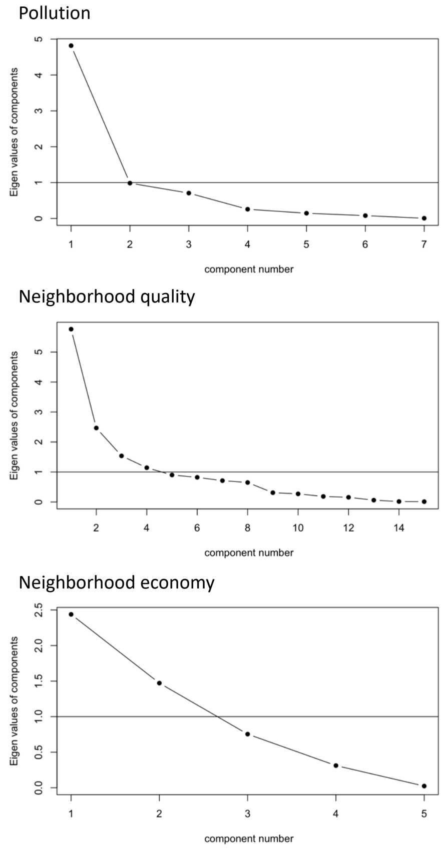


**Supplementary Figure 29.** Scree plots of environmental pollution, neighbourhood quality and neighbourhood economy measures.

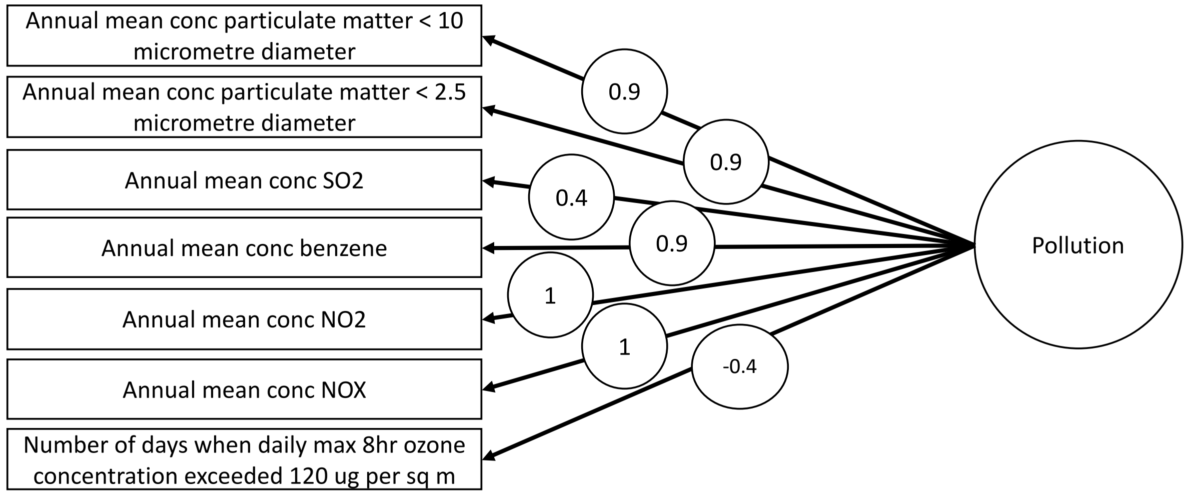


**Supplementary Figure 30.** Factor structure of environmental pollution measures.

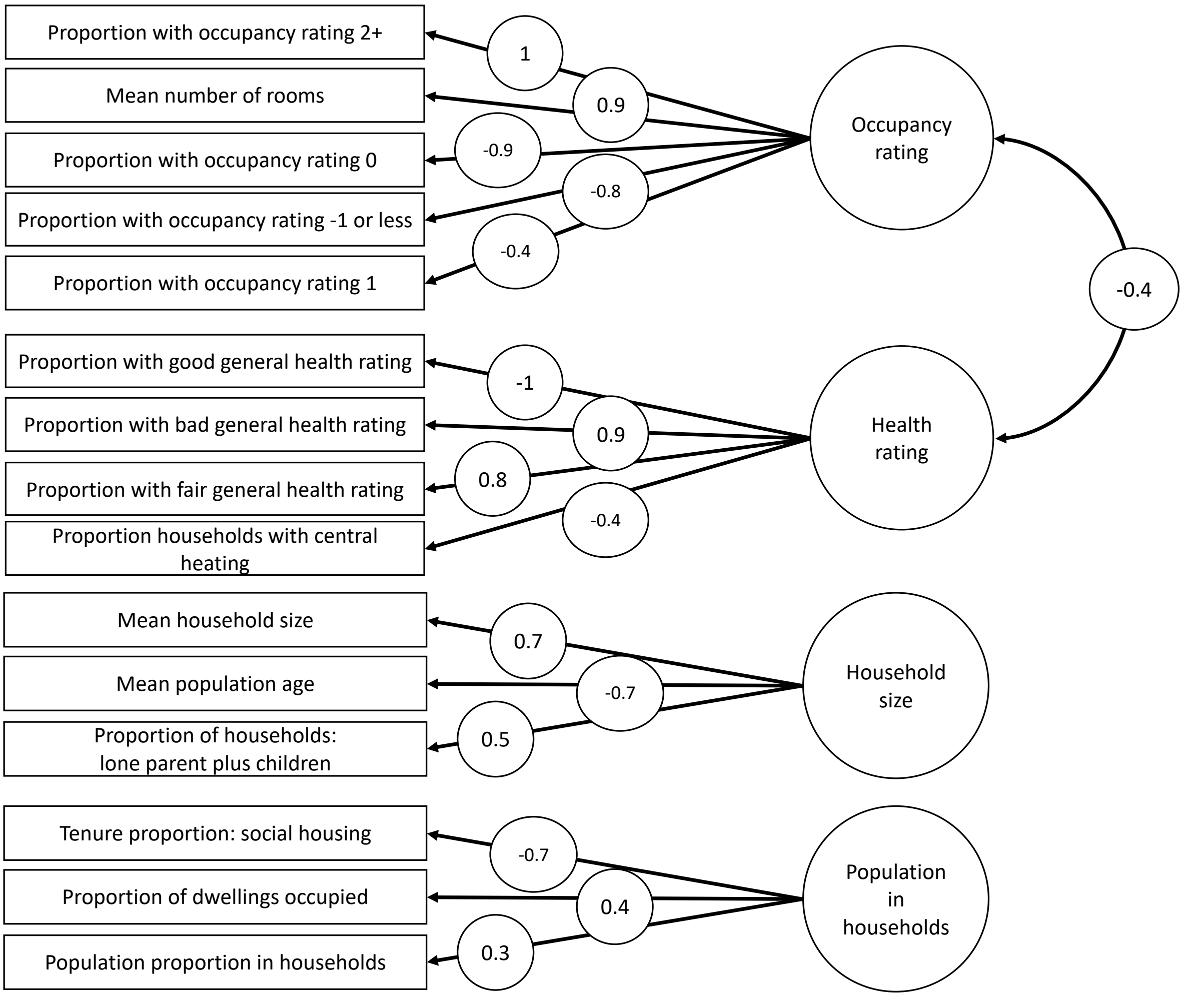


**Supplementary Figure 31.** Factor structure of neighbourhood quality measures.

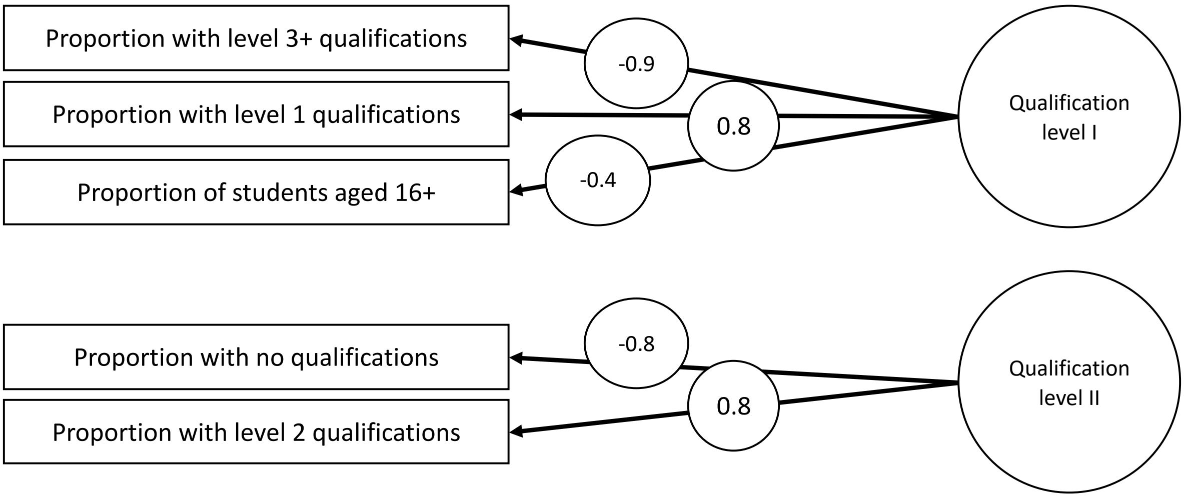


*Note.* Due to no theoretical coherence of qualification level II factor, only qualification level I factor was used in mediation analyses.

**Supplementary Figure 32.** Factor structure of neighbourhood economy measures.

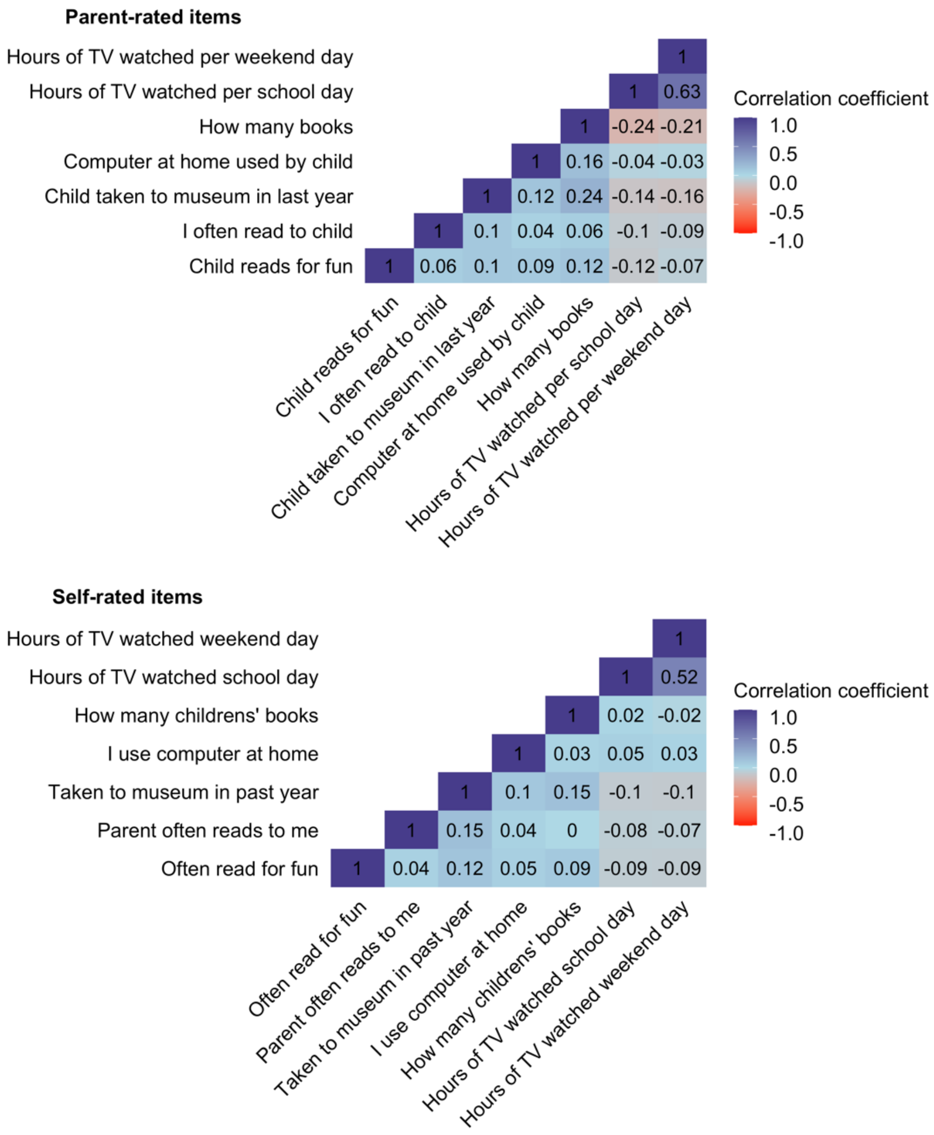


**Supplementary Figure 33.** Correlations between parent and self-rated home environment items at age 9.

Parent-rated items

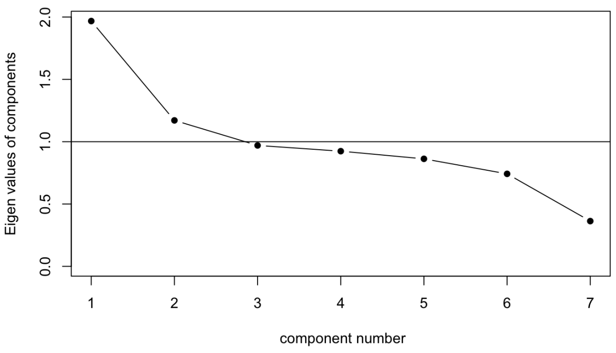


Self-rated items

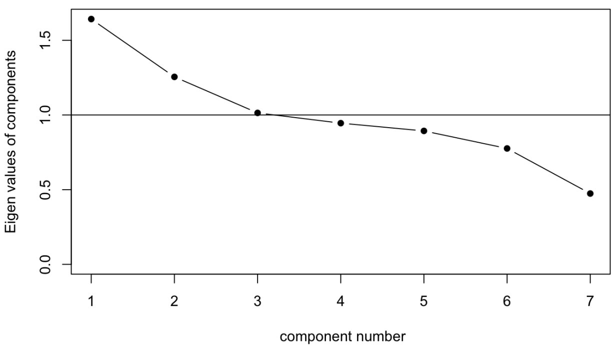


**Supplementary Figure 34.** Scree plots of parent and self-rated home environment items at age 9.

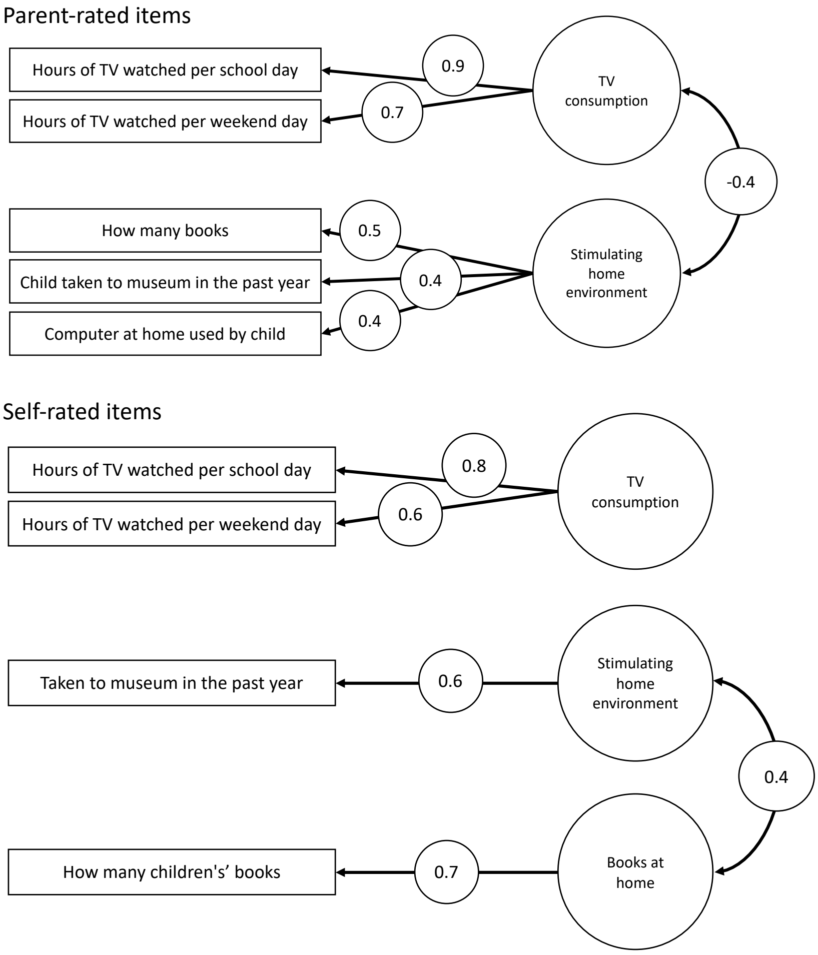


**Supplementary Figure 35.** Factor structure of parent and self-rated home environment items at age 9.

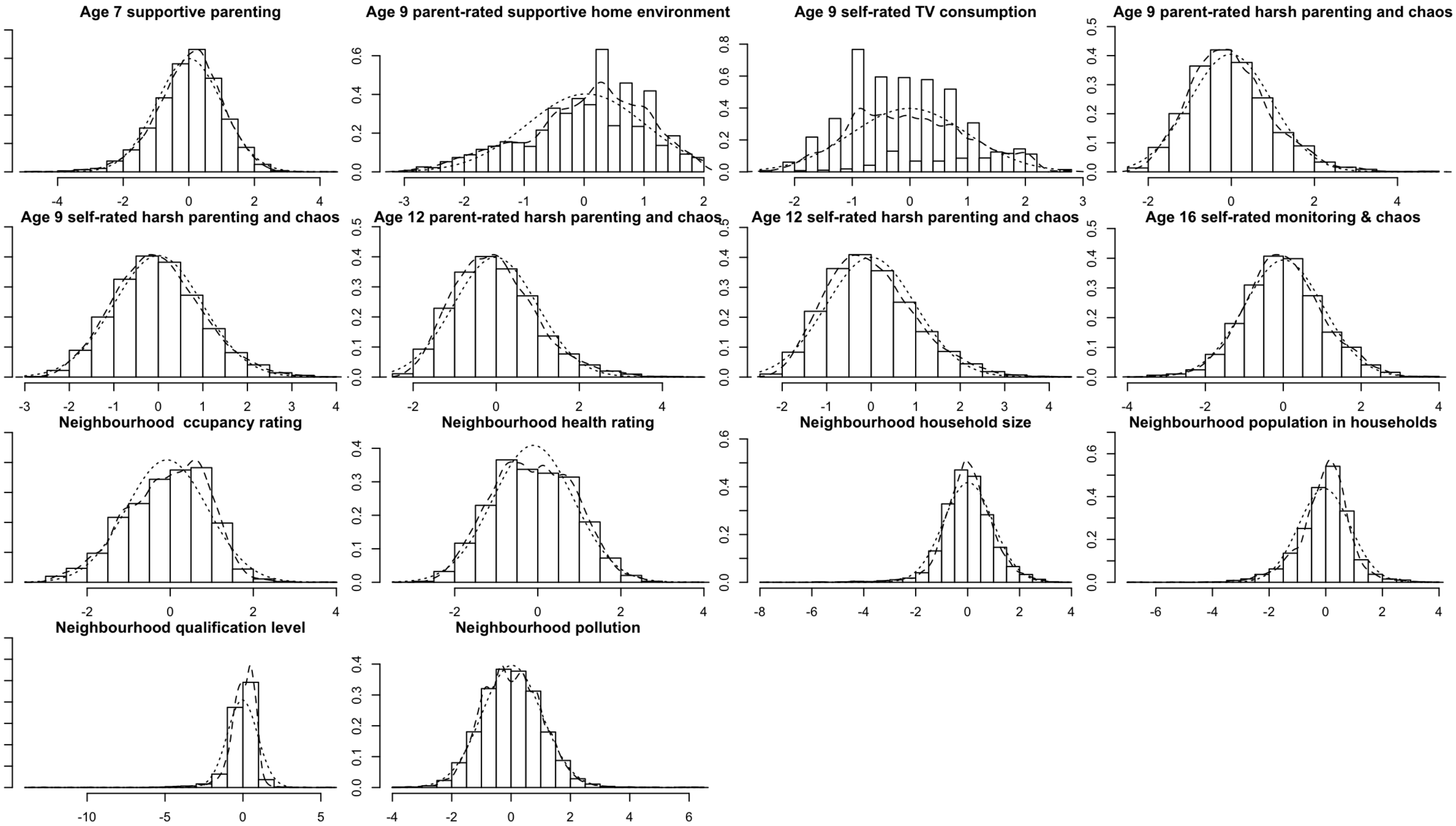


**Supplementary Figure 36.** Distributions of cross-sectional composites.

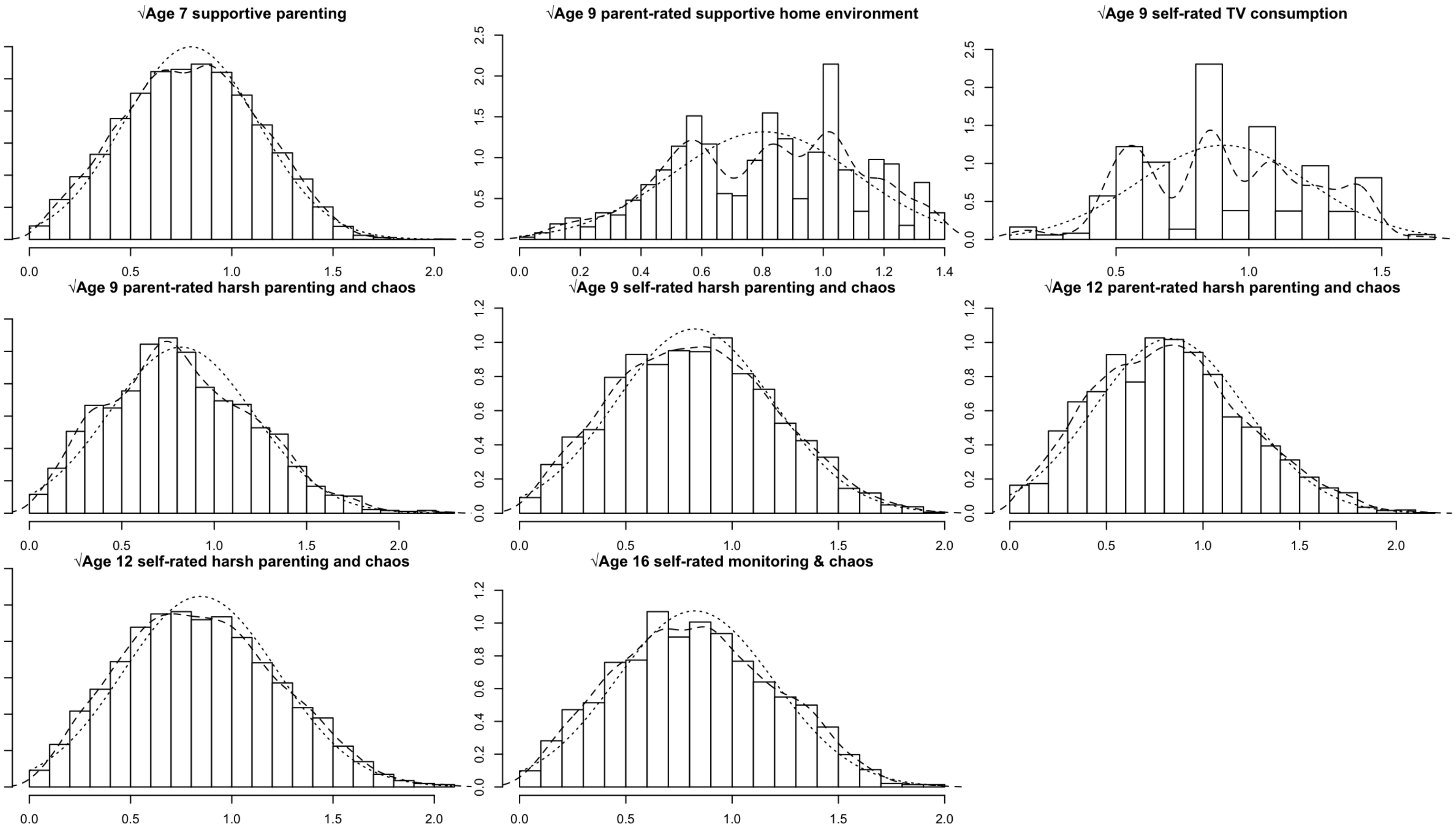


**Supplementary Figure 37.** Distributions of square root transformed cross-sectional composites.

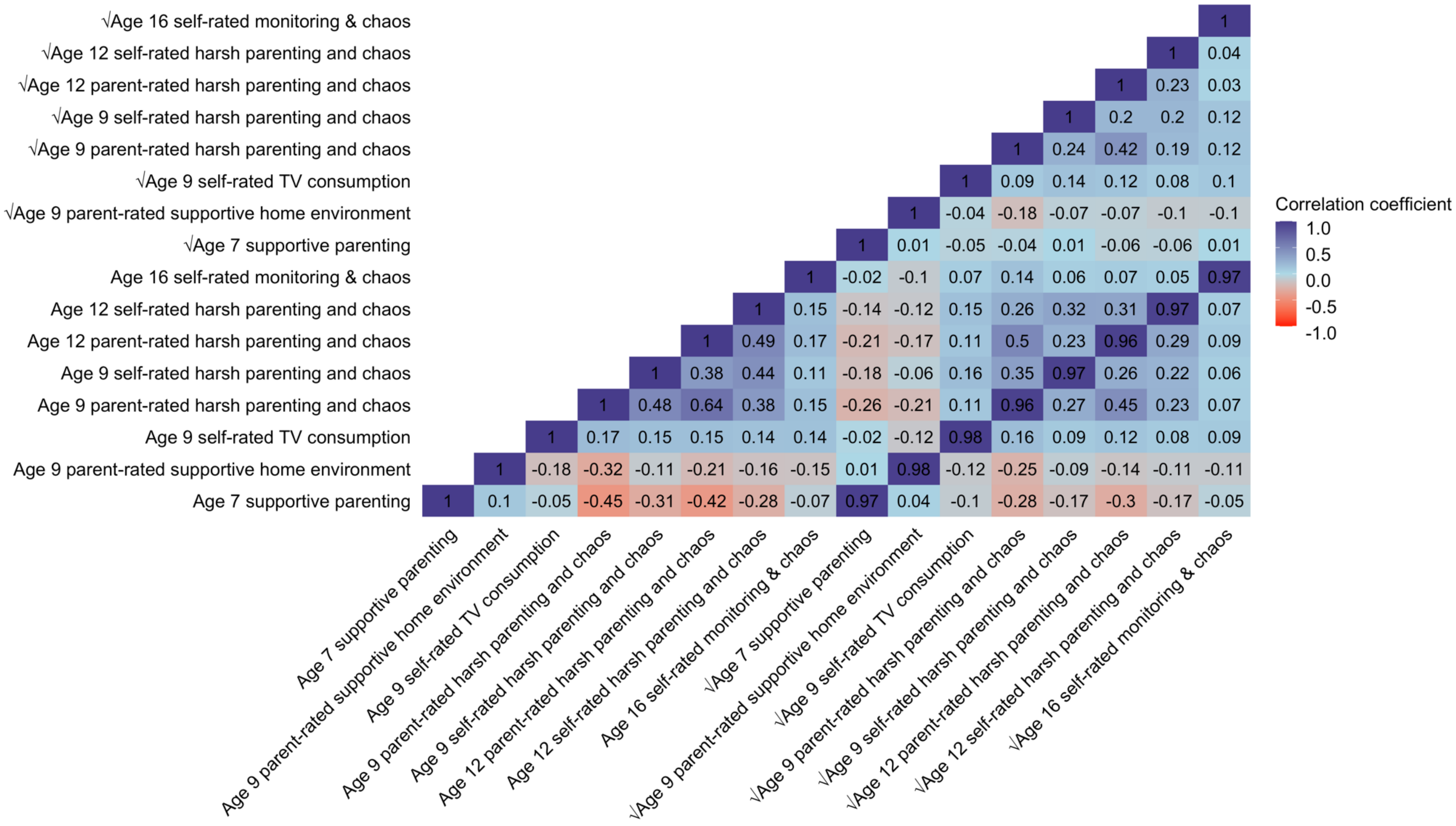


**Supplementary Figure 38.** Correlations between untransformed and square root transformed cross-sectional composites.

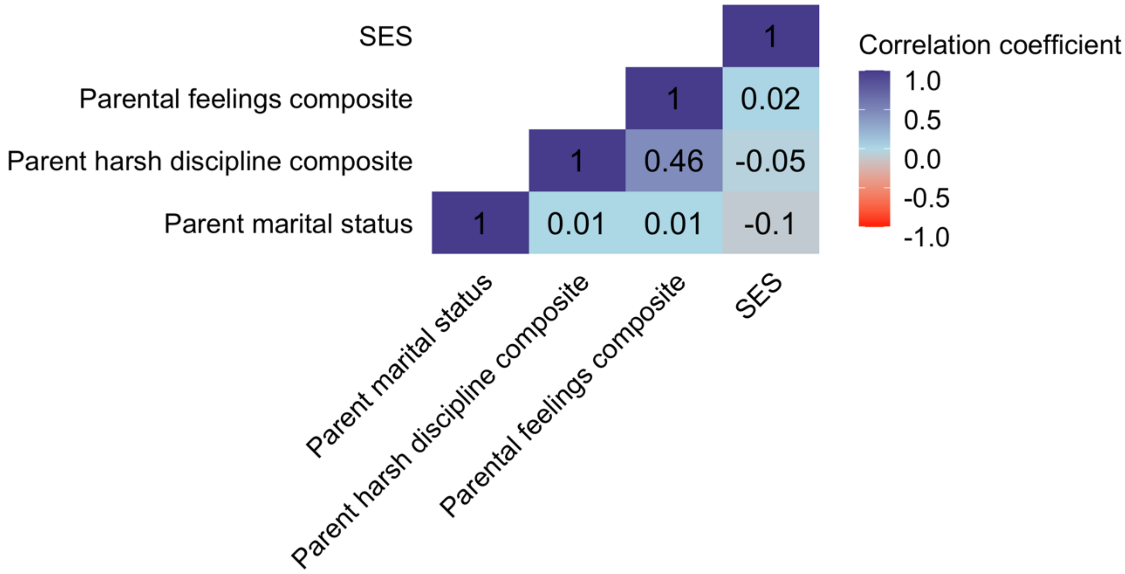


**Supplementary Figure 39.** Correlations between parent-rated environmental variables at age 7.

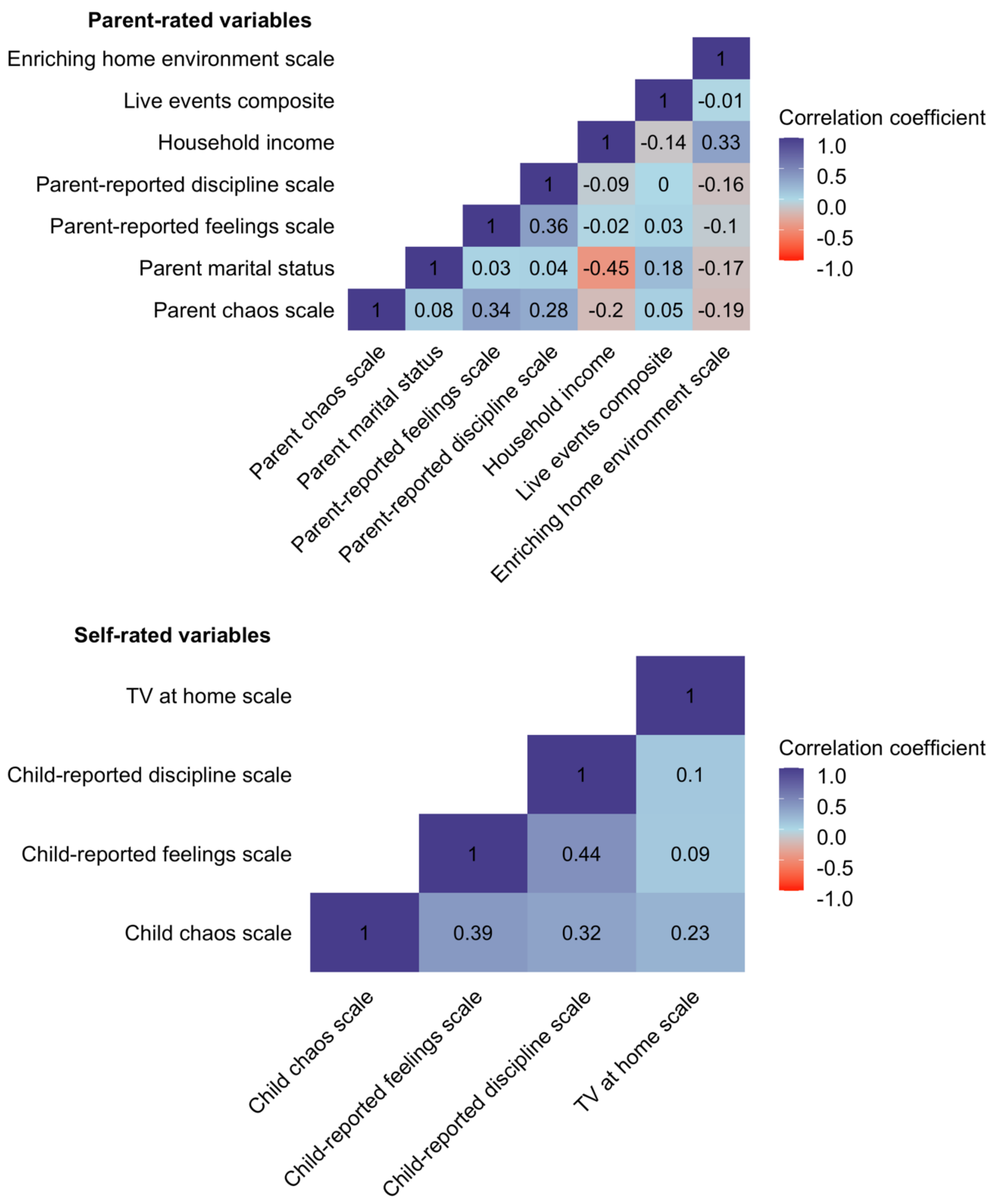


**Supplementary Figure 40**. Correlations between parent and self-rated environmental variables at age 9.

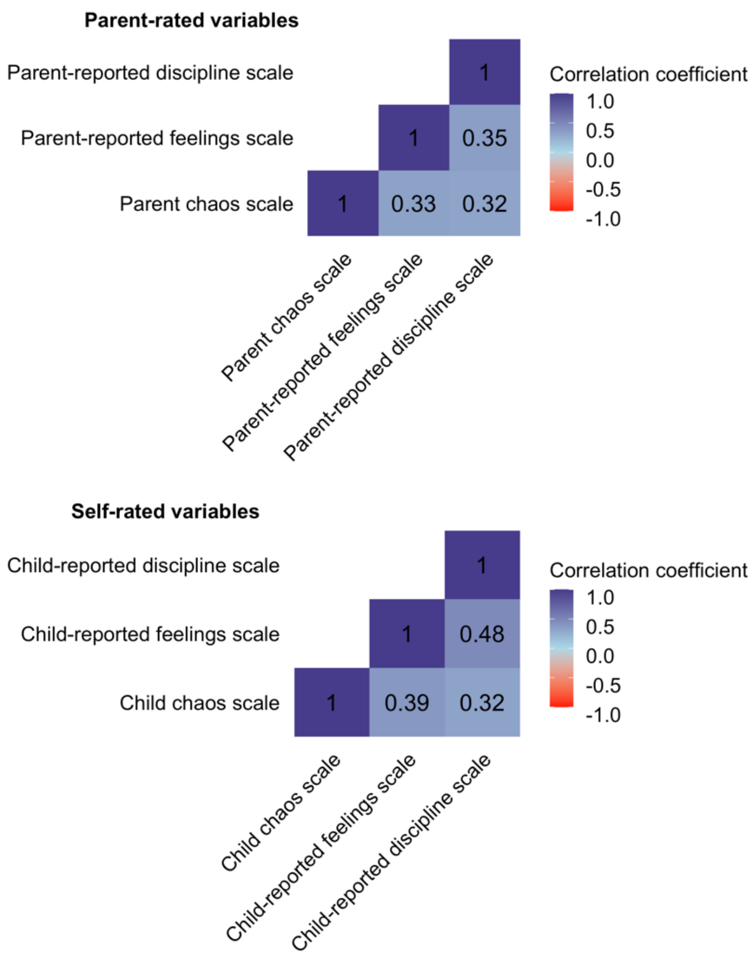


**Supplementary Figure 41.** Correlations between parent and self-rated environmental variables at age 12.

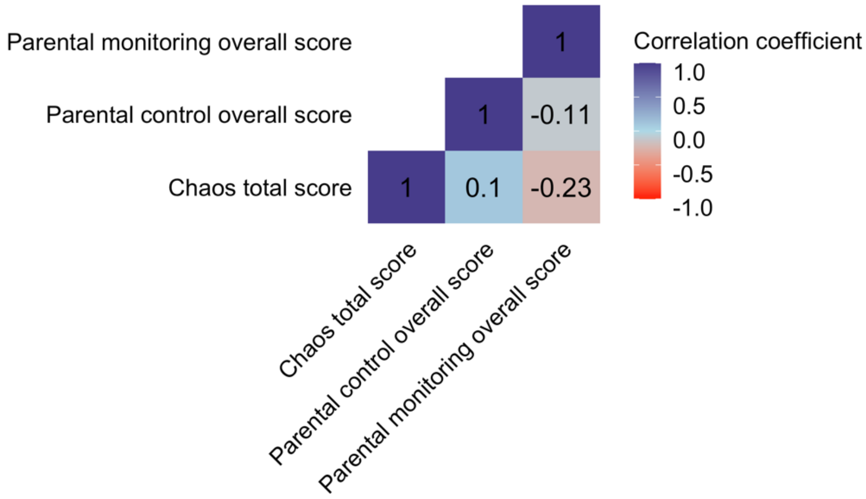


**Supplementary Figure 42.** Correlations between self-rated environmental variables at age 16.

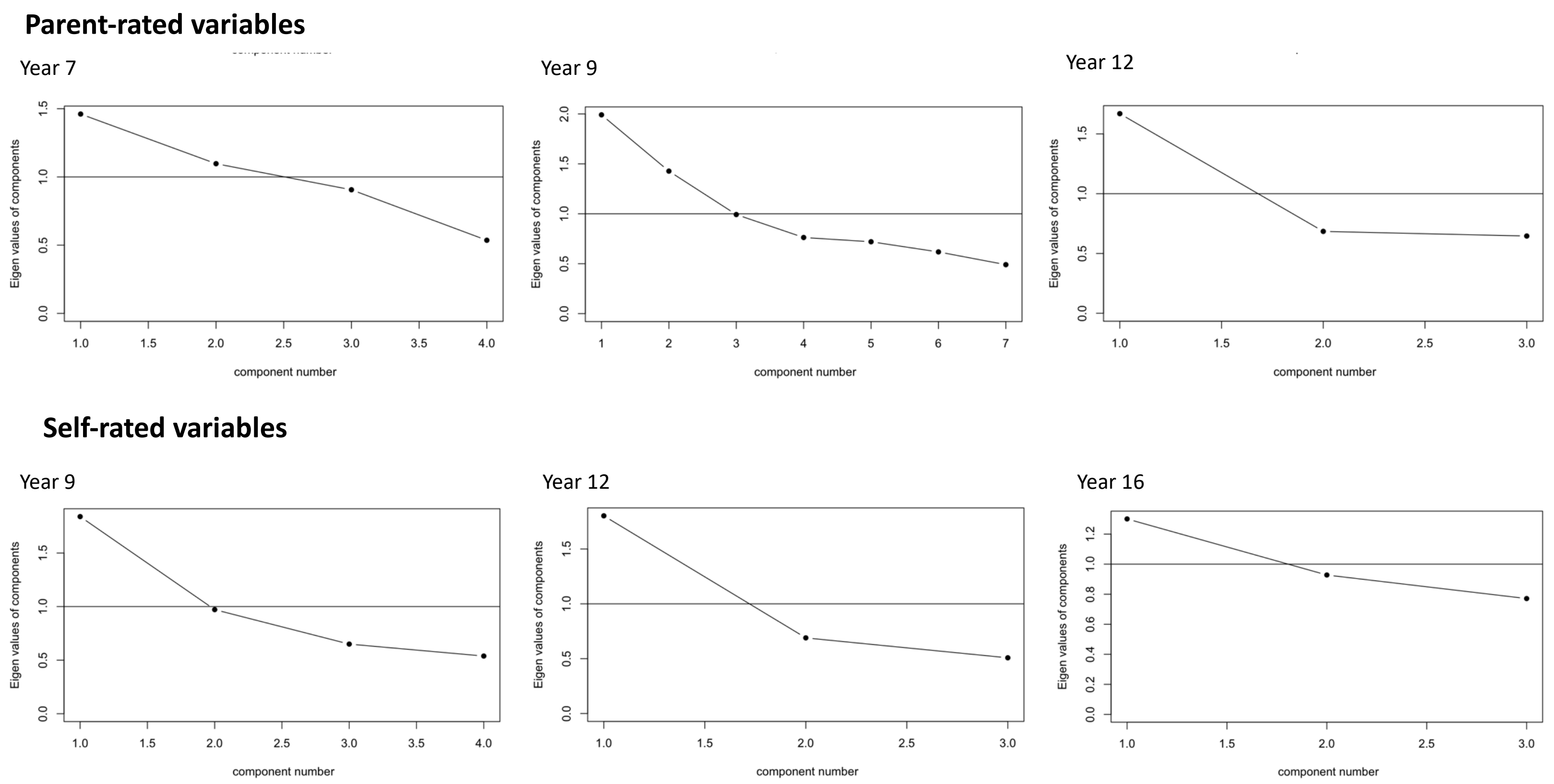


**Supplementary Figure 43.** Scree plots of parent and self-rated environmental variables at first contact and ages 7, 9, 12 and 16.

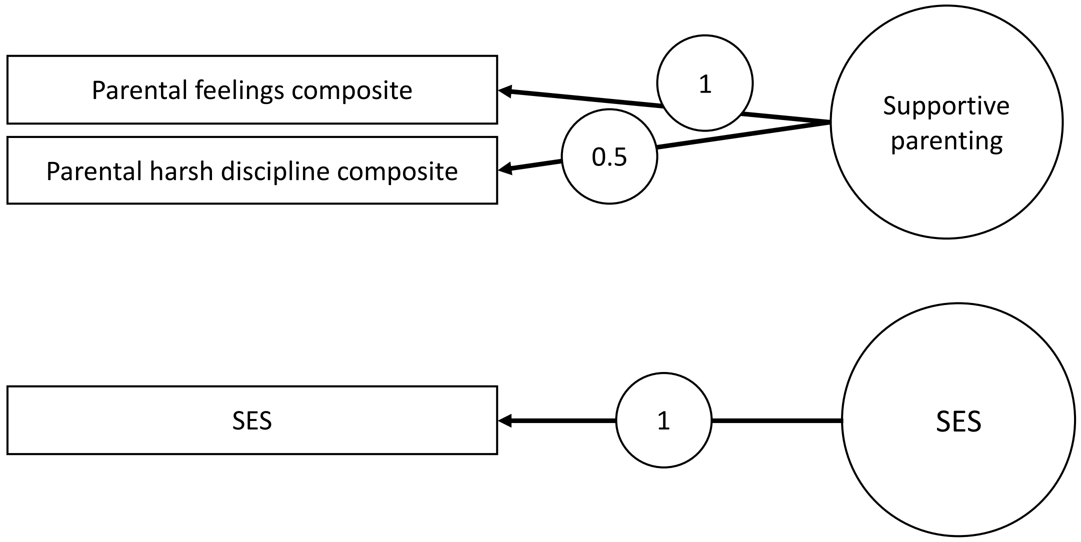


**Supplementary Figure 44.** Factor structure of parent-rated environmental variables at age 7.

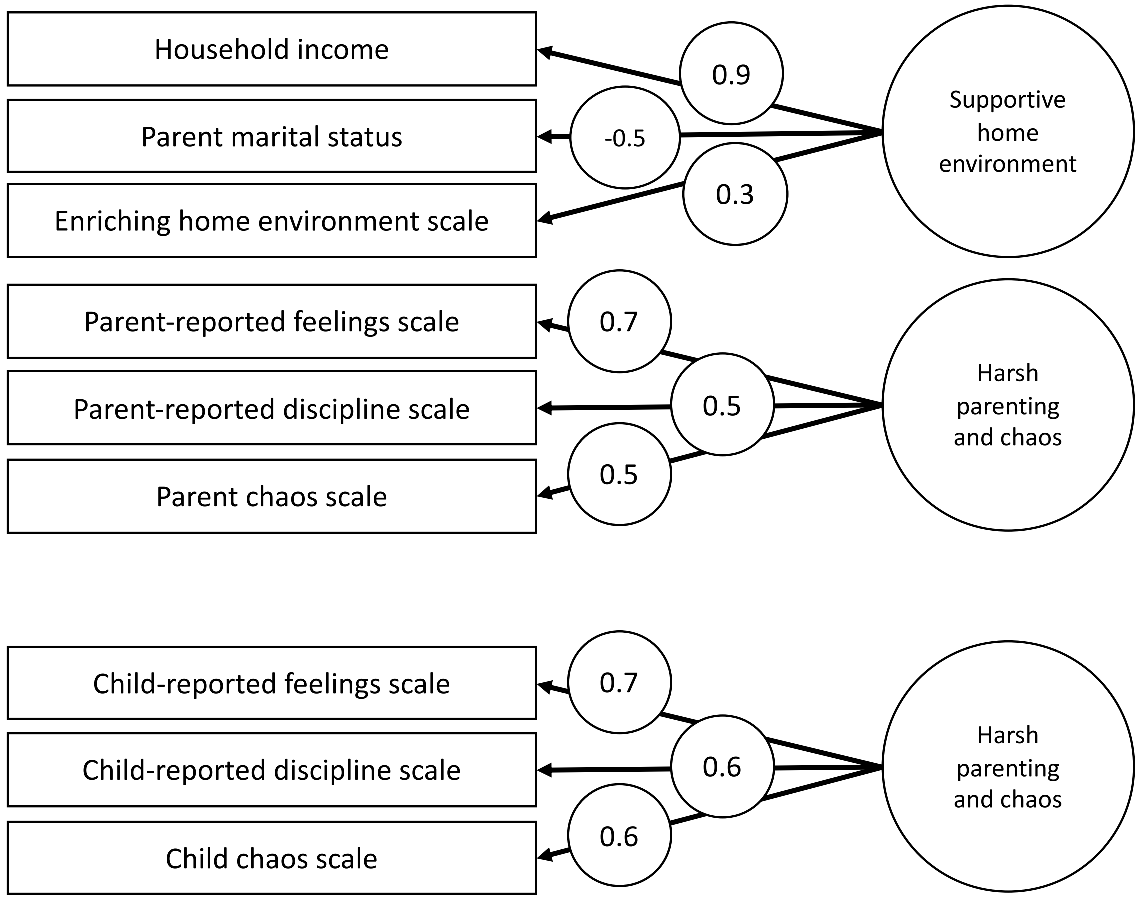


**Supplementary Figure 45.** Factor structure of parent and self-rated environmental variables at age 9.

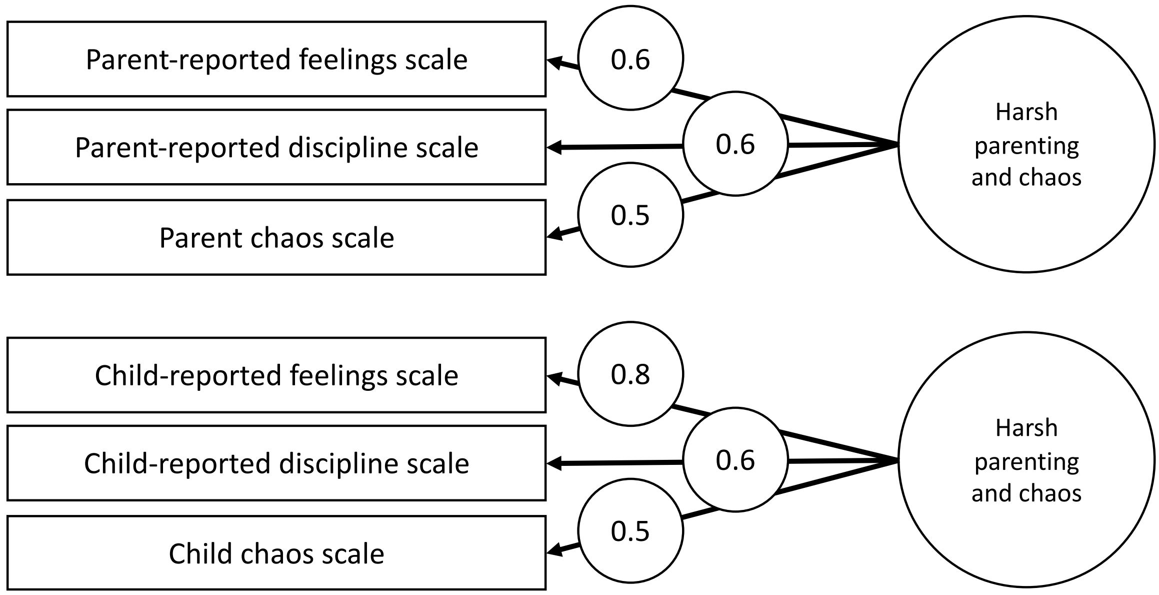


**Supplementary Figure 46.** Factor structure of parent and self-rated environmental variables at age 12.

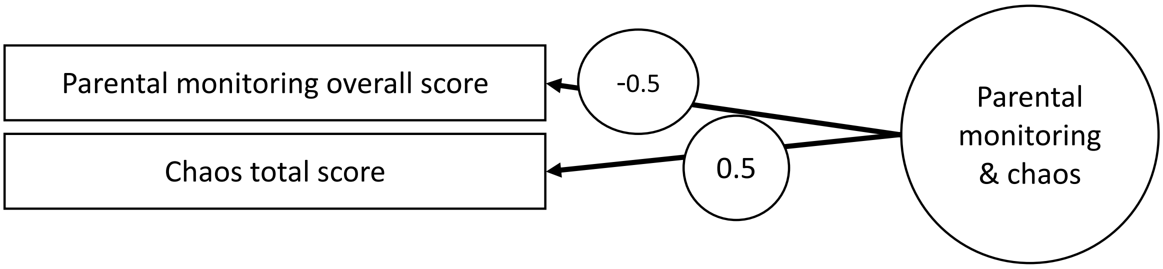


**Supplementary Figure 47.** Factor structure of self-rated environmental variables at age 16.

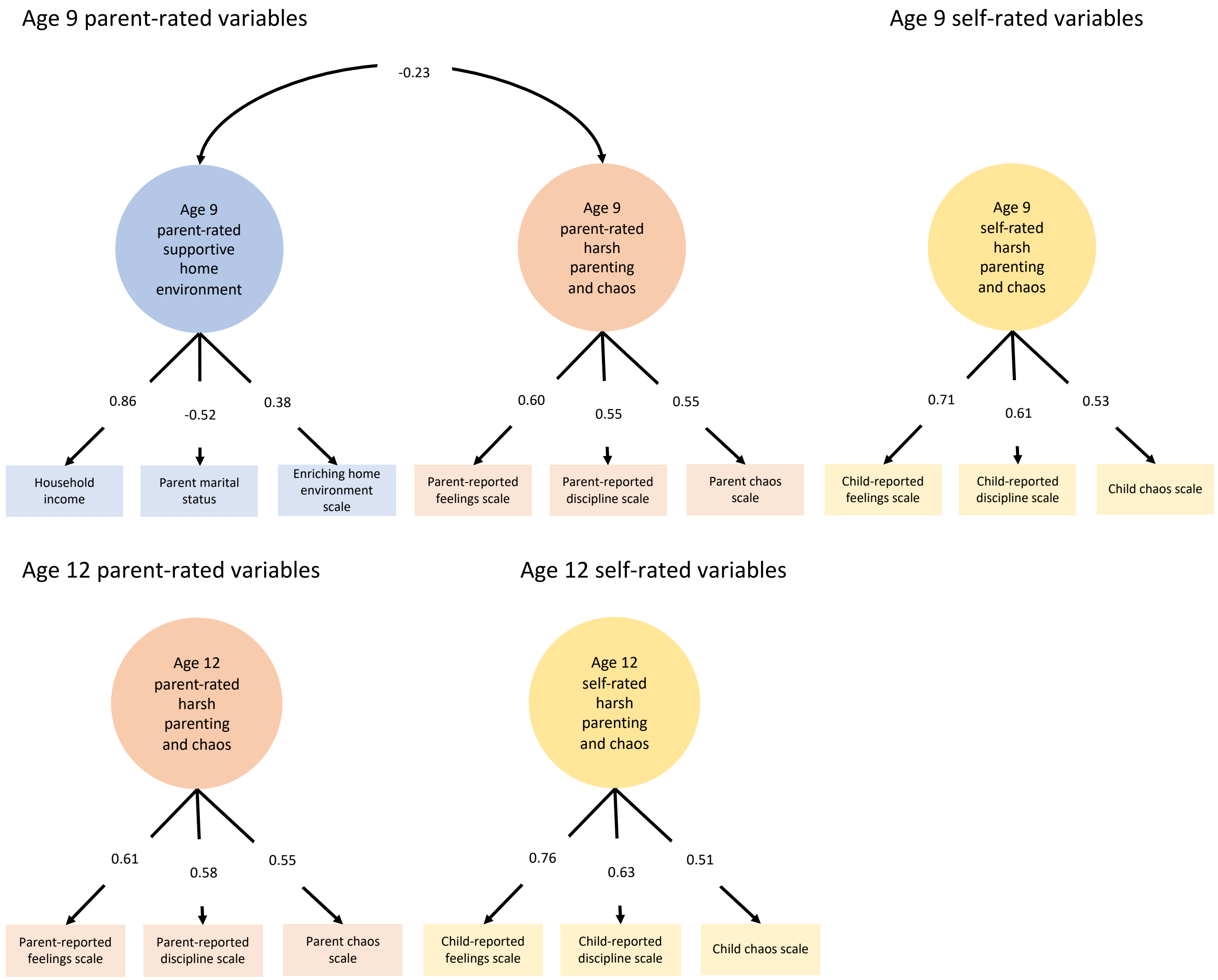


**Supplementary Figure 48.** CFA models of parent and self-rated latent environmental composites at ages 9 and 12.

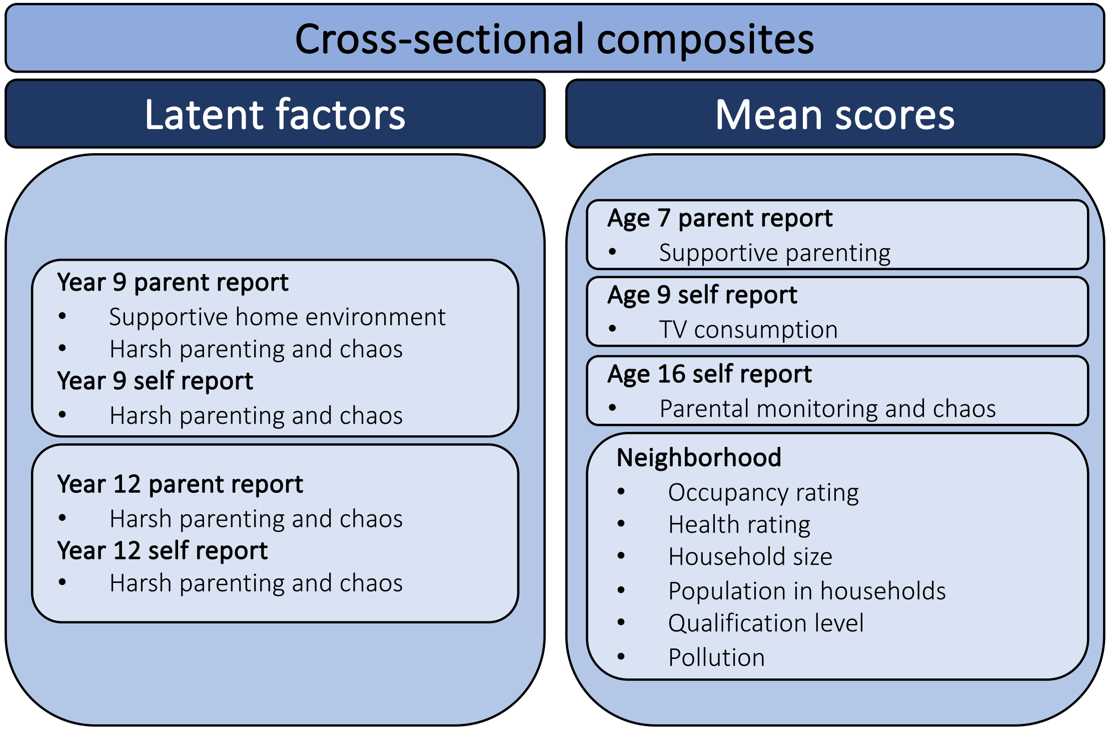


**Supplementary Figure 49.** Summary of the family environment composites.

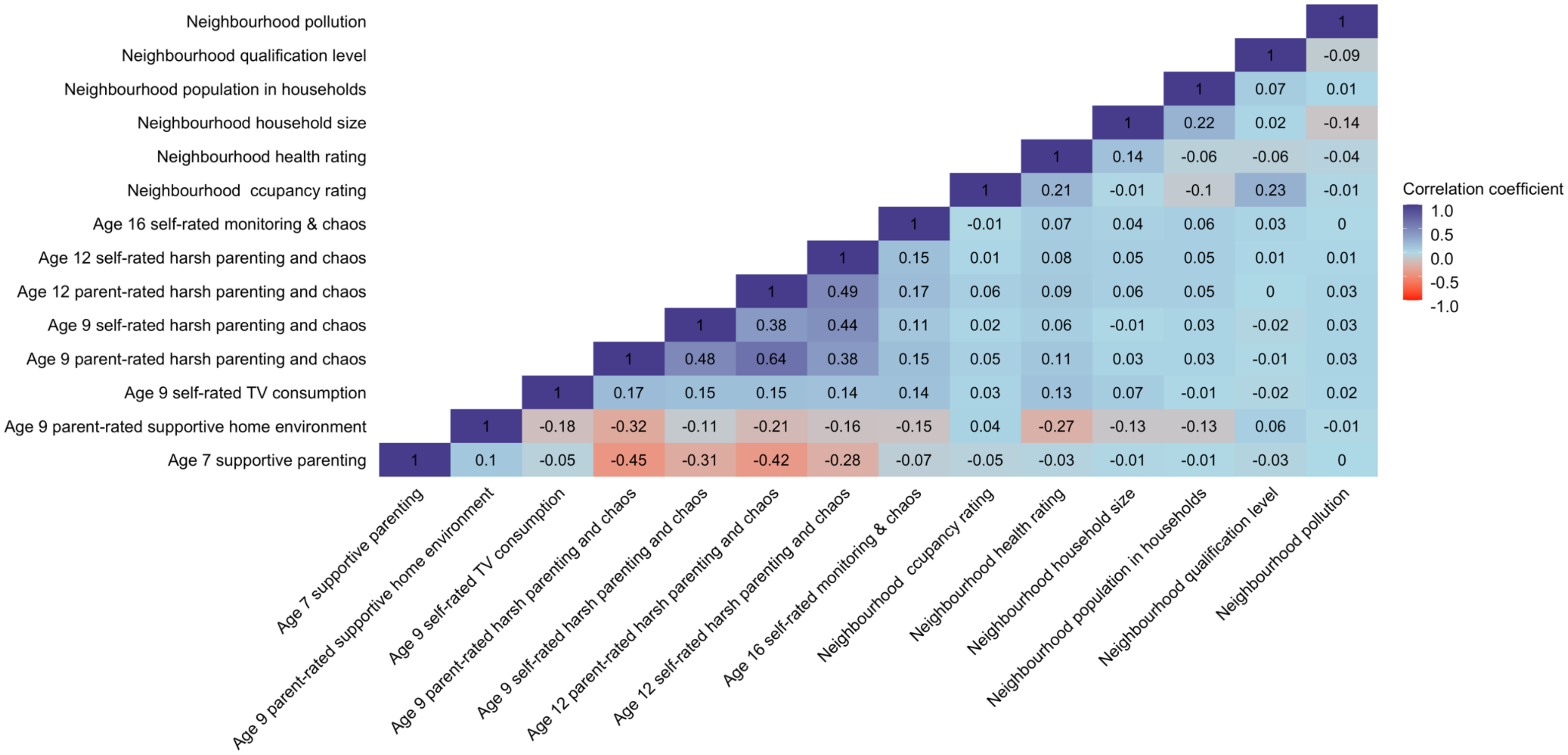


**Supplementary Figure 50.** Correlations between family environment composites.

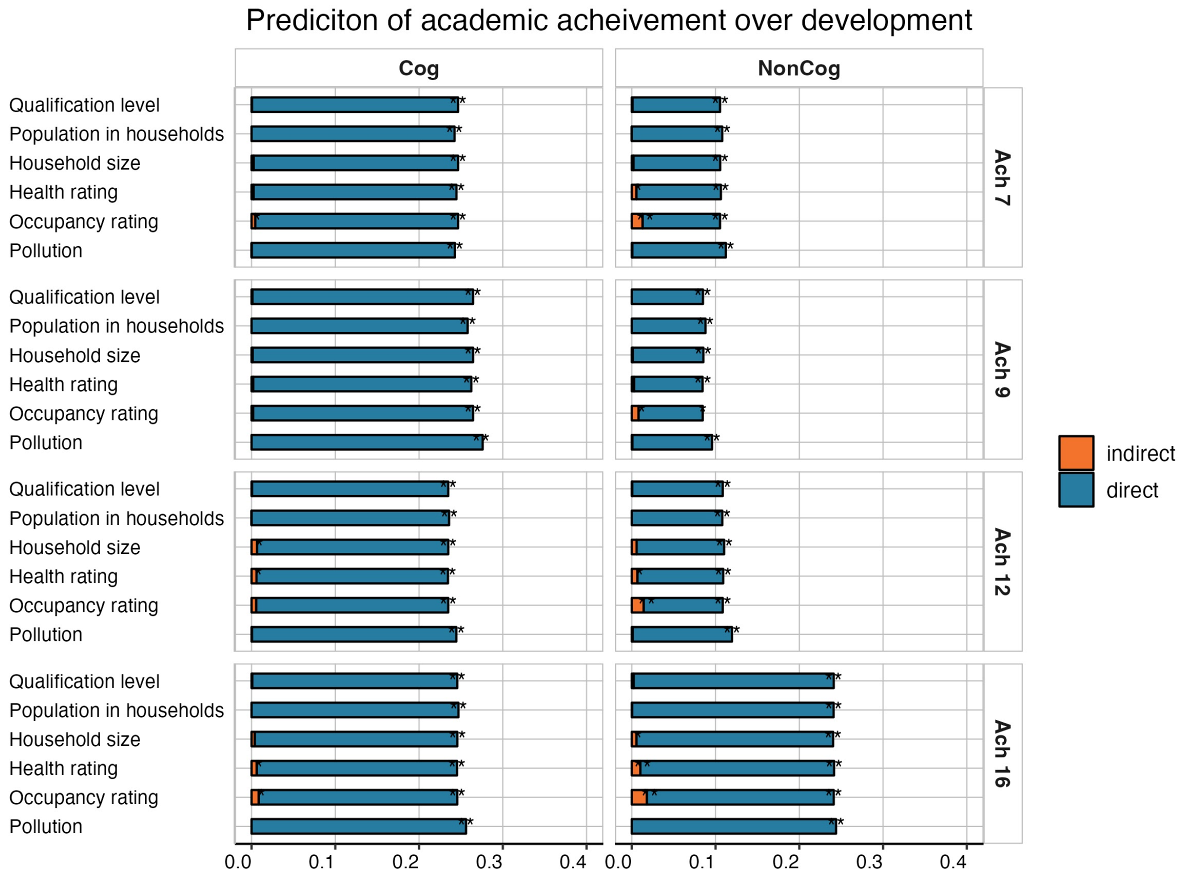


**Supplementary Figure 26.** Mediation models estimating the indirect effects of the neighbourhood environments on the cognitive (Cog) and noncognitive (NonCog) polygenic score prediction of academic achievement over development

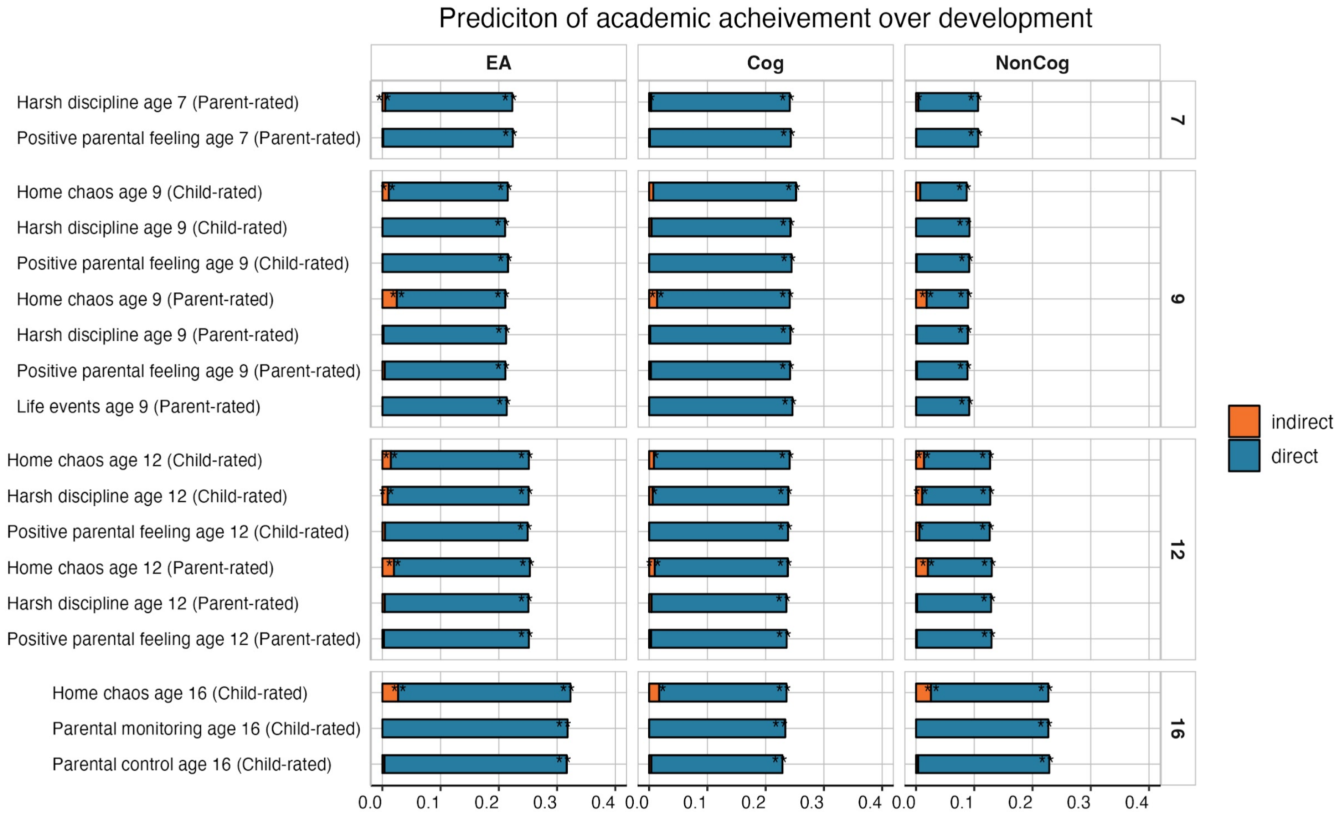


**Supplementary Figure 27.** Mediation models estimating the indirect effects of individual family environment measures on the educational attainment (EA), cognitive (Cog) and noncognitive (NonCog) polygenic score prediction of academic achievement over development.

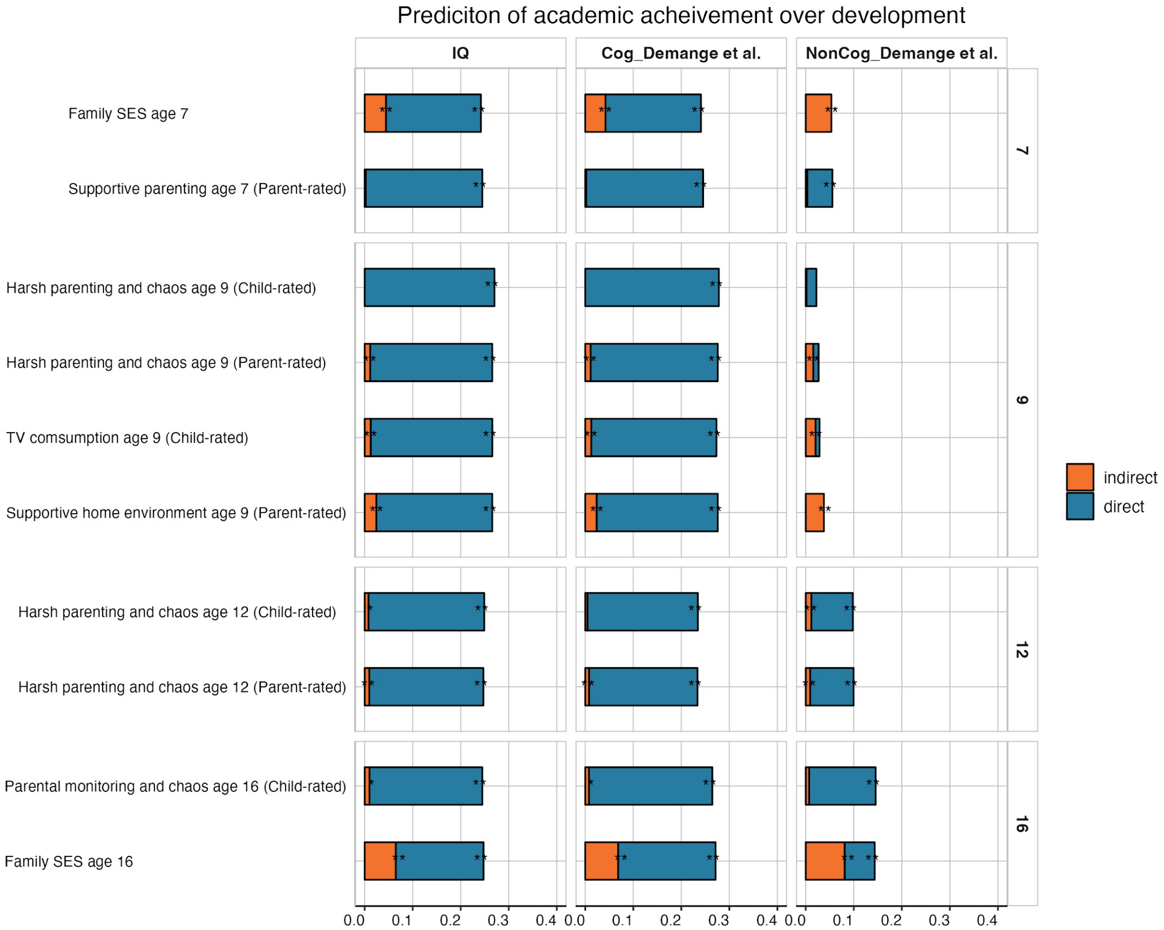


**Supplementary Figure 28.** Mediation models estimating the indirect effects of the family environment composites on the intelligence (IQ), Cognitive (Cog_Demange et al.) and noncognitive (NonCog_Demange et al. ) polygenic score predictions of academic achievement over development. These polygenic scores were estimated using a different model to quantify cognitive and noncognitive polygenic scores, published by Demange et al., 2021 (4).

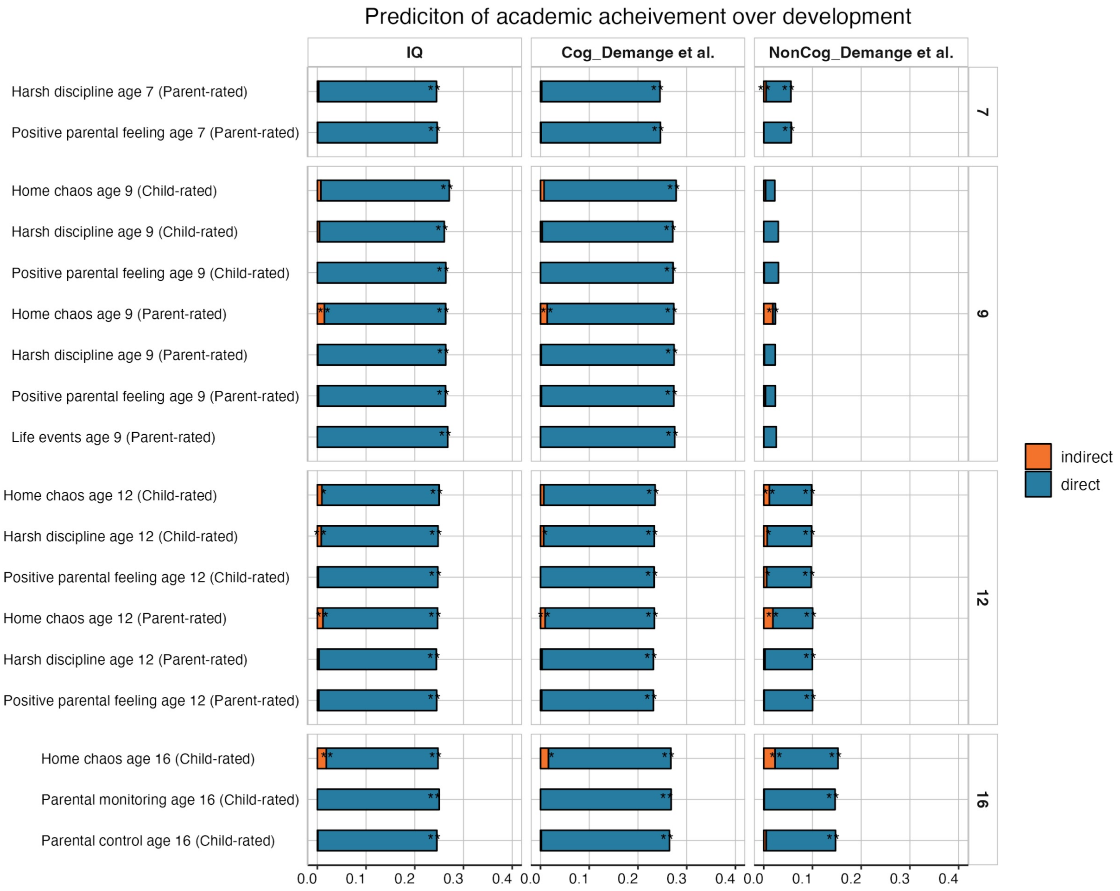


**Supplementary Figure 29.** Mediation models estimating the indirect effects of individual family environment measures on the intelligence (IQ), Cognitive (Cog_Demange et al.) and noncognitive (NonCog_Demange et al.) polygenic score predictions of academic achievement over development. These polygenic scores were estimated using a different model to quantify cognitive and noncognitive polygenic scores, published by Demange et al., 2021 (4).

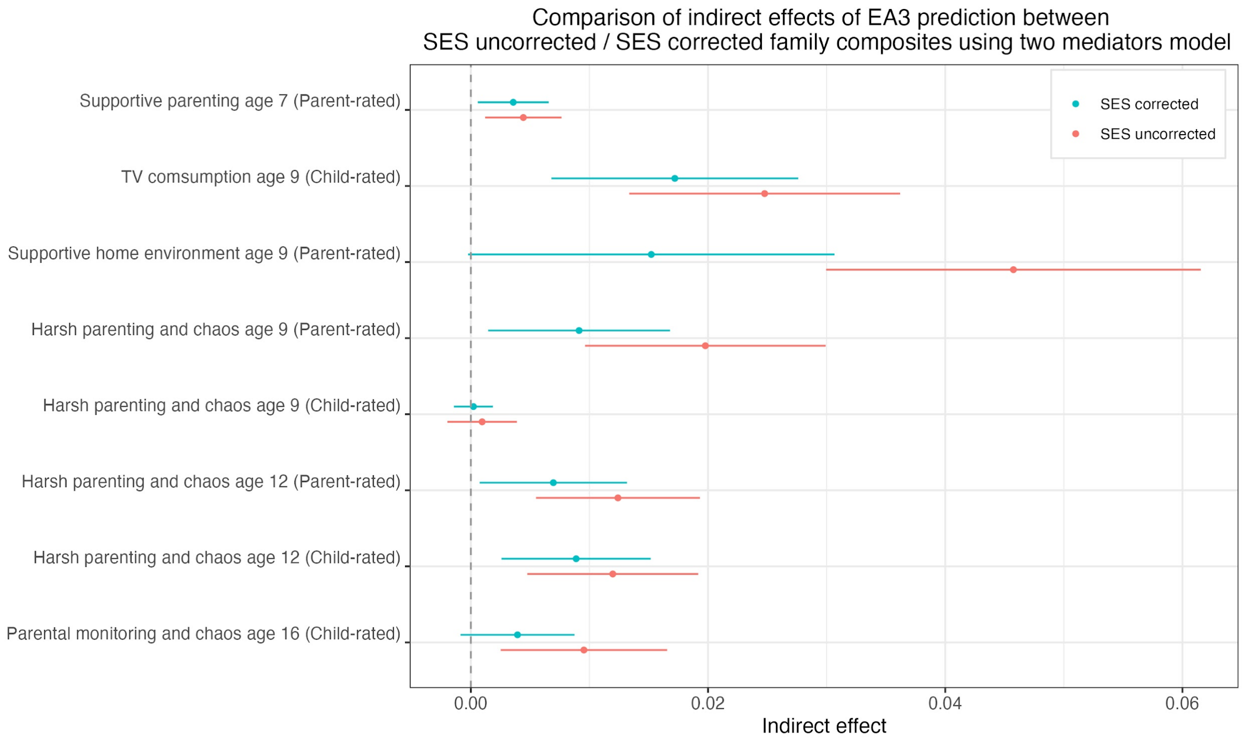


**Supplementary Figure 30.** Comparison of indirect effects of the educational attainment (EA3) polygenic score on academic achievement over development before and after accounting for socioeconomic status using a series of two-mediators model.

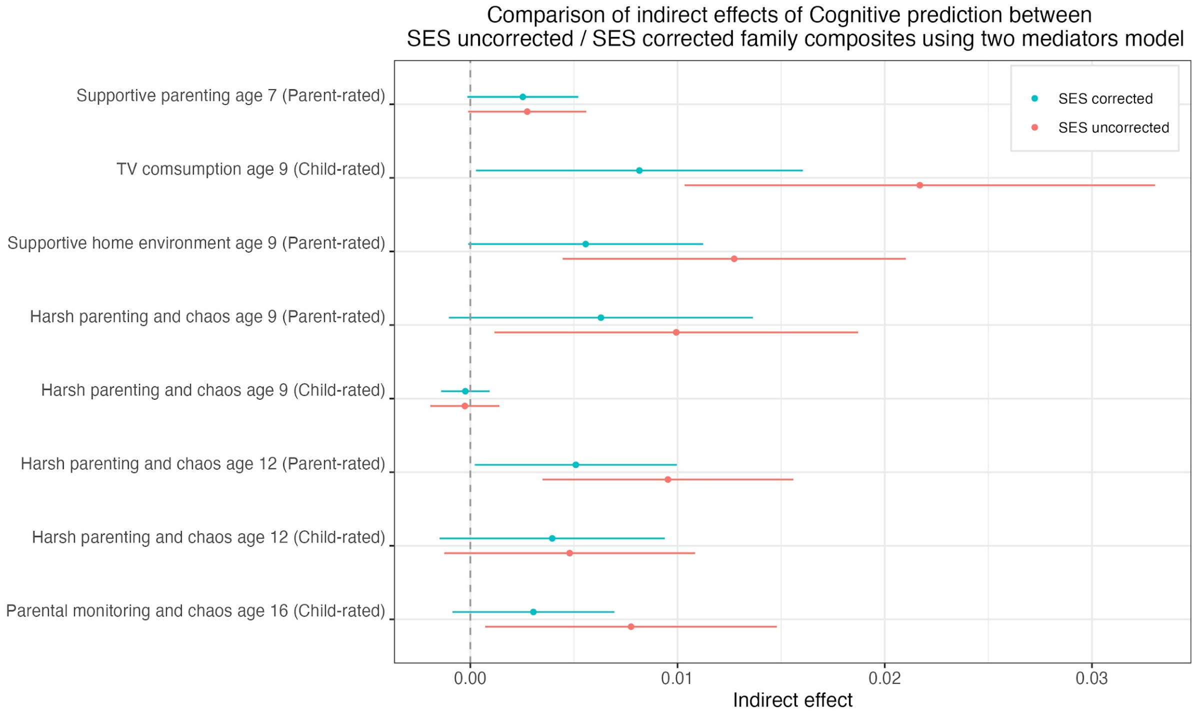


**Supplementary Figure 31.** Comparison of indirect effects of the cognitive polygenic score on academic achievement over development before and after accounting for socioeconomic status using a series of two-mediators model.

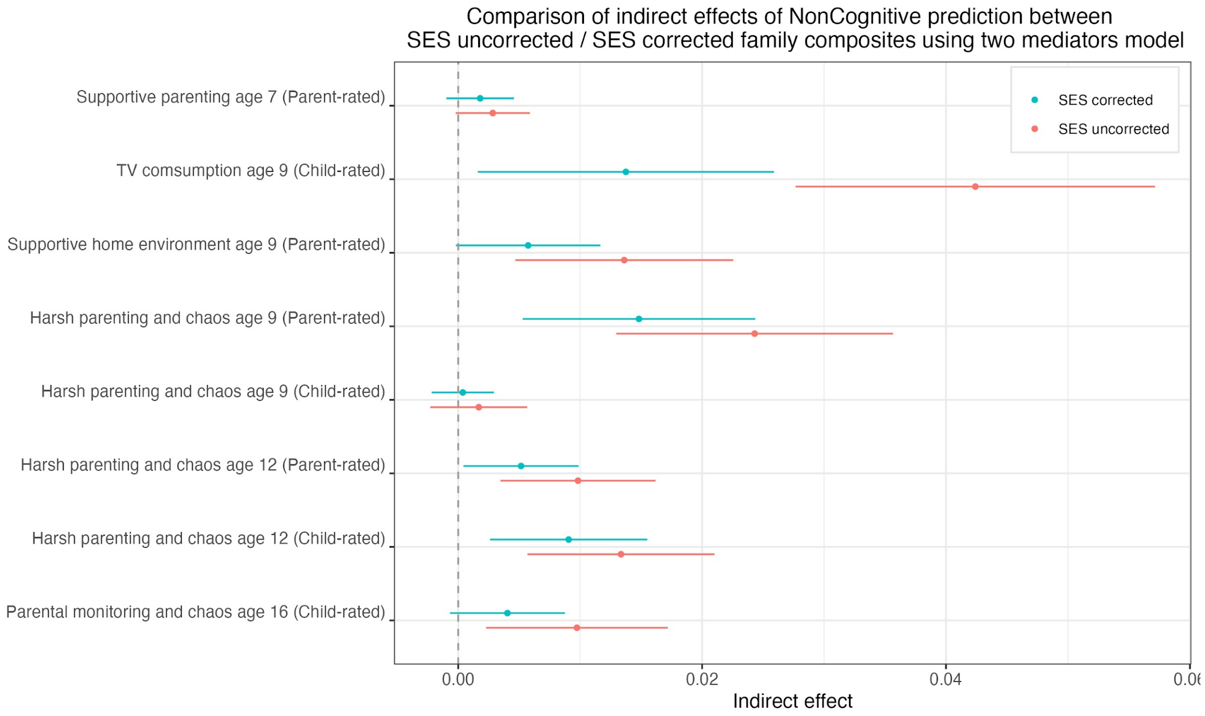


**Supplementary Figure 32.** Comparison of indirect effects of the noncognitive polygenic score on academic achievement over development before and after accounting for socioeconomic status using a series of two-mediators model.

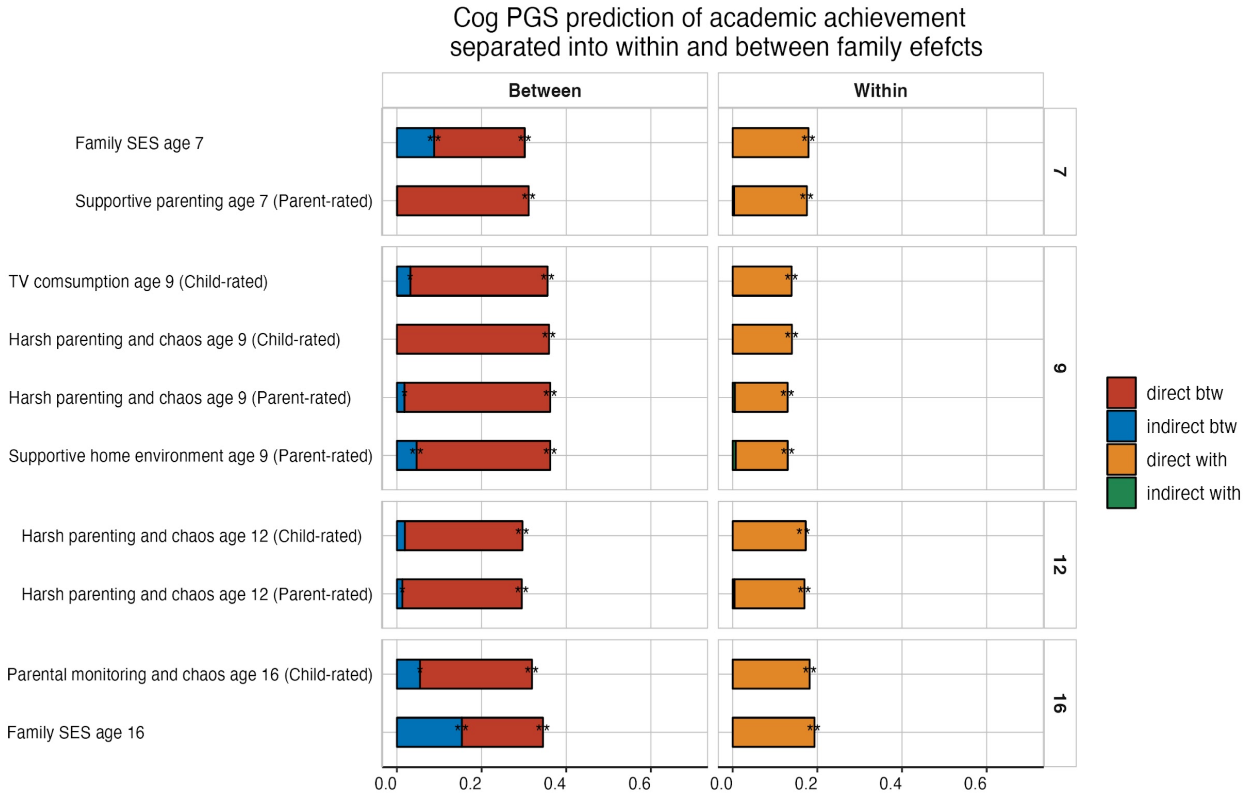


**Supplementary Figure 33:** Mediation models estimating the indirect effects of family environment composite measures on the cognitive (Cog) polygenic score predictions of academic achievement over development, separated into within and between family effects.

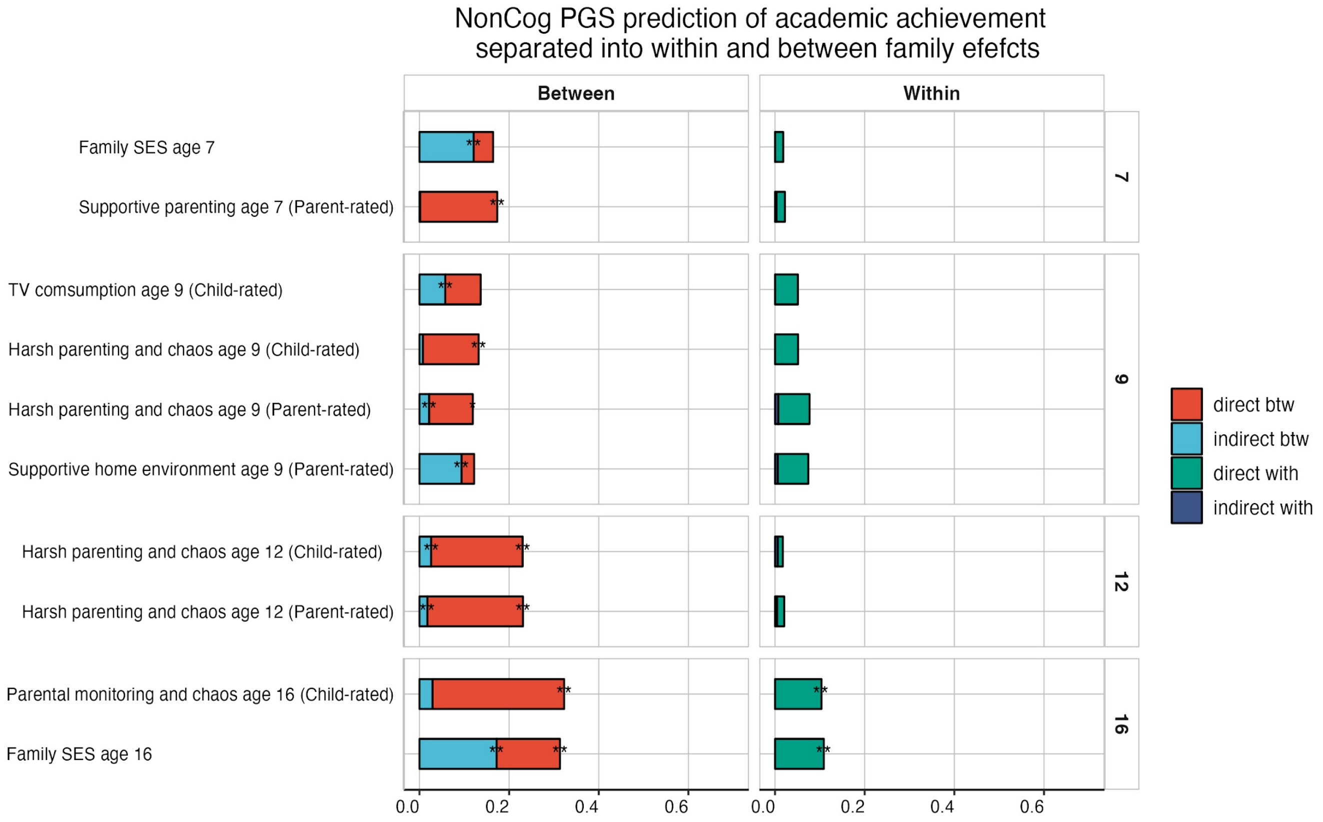


**Supplementary Figure 34:** Mediation models estimating the indirect effects of family environment composite measures on the noncognitive (NonCog) polygenic score predictions of academic achievement over development, separated into within and between family effects.
